# Spatially localized fMRI metrics as predictive and highly distinct state-independent fingerprints

**DOI:** 10.1101/2021.08.03.454862

**Authors:** Daouia I. Larabi, Martin Gell, Enrico Amico, Simon B. Eickhoff, Kaustubh R. Patil

## Abstract

Precision medicine and the investigation of brain-behavior associations require biomarkers that are stable (low intraindividual variability) and unique (high interindividual variability) at the same time, hence calling them “fingerprints”. The functional connectome (FC) has good “fingerprint properties”, as individuals can be accurately identified in a database based on their FC. Importantly, research has shown lower intraindividual variability of more localized measures of brain function such as regional homogeneity (ReHo) and (fractional) amplitude of low-frequency fluctuations ((f)ALFF), compared to the FC. Here, with fMRI data from two publicly available datasets we demonstrate that individuals can be identified with near-perfect accuracies using local functional fingerprints, and especially the regional homogeneity (ReHo) fingerprint. Further analyses reveal that the dorsal attention network contributes most to the individual “uniqueness” of the ReHo fingerprint. Moreover, using a machine-learning setup, we show that the small intraindividual ReHo fingerprint variability across sessions is meaningful for explaining individual-level intelligence. Last, using two other publicly available datasets, clinical applicability is shown with high fingerprint accuracies and a significant correlation between fingerprint stability and intelligence in individuals with schizophrenia. Altogether, our findings suggest that the ReHo fingerprint is a good candidate for further exploration of applicability in precision medicine.

## Introduction

Large variability across individuals (i.e., interindividual variability) is seen in phenotypic traits, including physical traits and cognitive abilities, as well as brain organization. Identifying the interindividual variability in functional brain organization that is predictive of cognitive and clinical properties at the individual level is important for developing viable biomarkers for precision medicine and *precision psychiatry* specifically (Fernandes et al., 2017). Precision psychiatry is the tailoring of treatment to the individual. It requires trait-like representations of brain function that are simultaneously reliable (i.e., show low *intraindividual* variability) and unique (i.e., show high *interindividual* variability), i.e., representations that are stable within an individual but vary across individuals. Intraindividual stability independent of an individual’s current state is a prerequisite for clinical application, as biomarkers should reflect underlying traits or disease and not the current state. Representations should also be unique for an individual, as the interindividual variability in these representations can then be utilized for the prediction of individual-level behavior with machine learning algorithms (Chen et al., 2020). Thus, a trait-like state-independent neural fingerprint that captures interindividual differences can shed light on fundamental brain organization as well as facilitate translational applications.

Until now, the functional connectome (FC) has been *the* choice to predict cognition and behavior. It reflects whole-brain patterns of synchrony in fluctuations of the blood-oxygen-level-dependent (BOLD) signal, as measured with functional magnetic resonance imaging (fMRI). The FC is usually calculated as Pearson’s correlations between the time courses of all possible pairs of regions. It has been termed a functional connectome “fingerprint” as it is highly stable and sufficiently unique to identify an individual across sessions (Finn et al., 2015; Seitzman et al., 2019). The FC has also been shown to capture interindividual differences (Finn and Rosenberg, 2021; Nostro et al., 2018): the variability in the FC fingerprint has been associated with development, aging, intelligence, and psychiatric disorders (Amico and Goñi, 2018; Kaufmann et al., 2018, 2017; Miranda-Dominguez et al., 2014; Noble et al., 2019). More specifically, it can be indicative of psychiatric disorders such as schizophrenia, depression, or attention deficit disorder (Kaufmann et al., 2018). Hence, it fulfills the previously mentioned properties of low intraindividual variability and high interindividual variability subsequently called “fingerprint properties” in this manuscript.

However, even though the FC is most commonly used for the prediction of cognitive and clinical properties, several studies have shown lower test-retest reliability of global measures of brain function as opposed to local measures such as regional homogeneity (ReHo), amplitude of low-frequency fluctuations (ALFF), and fractional ALFF (fALFF) (Chen et al., 2015; Holiga et al., 2018; Jiang and Zuo, 2016). Specifically, local metrics show lower intraindividual variability, suggesting that these metrics are less dynamic than global metrics (Chen et al., 2015). Therefore, here, we compared the FC baseline “fingerprint” to local functional “fingerprints” within predefined brain regions based on the 264 Power coordinates (Power et al., 2011). We quantified fingerprint properties within the identification framework in which identification is the ability to identify an individual in a new database. Using the resting-state and task fMRI data of the Human Connectome Project (HCP) and four different ways to quantify identification, including the novel “ranking accuracy” and “forensic identification accuracy”, we found near-perfect identification accuracies when using ReHo, followed by ALFF, fALFF, and the FC. We extensively checked for replicability of these findings across various parcellations and resilience against confounding effects. Next, four follow-up analyses were conducted. First, using a novel optimization formulation, we investigated which regions contributed most to making the ReHo fingerprint “unique” at the individual level. Second, the previously mentioned fingerprint properties of reliability and idiosyncrasy are necessary but not *sufficient* for clinical applicability, given that fingerprints should also capture interindividual variance related to actual behavior (Finn and Rosenberg, 2021; Mantwill et al., 2021). We tested this by using both the fingerprints themselves and the *fingerprint stability* - i.e., their invariance across sessions - to predict crystallized and fluid intelligence in a machine-learning setup. This allowed us to test whether the (small) intraindividual fingerprint variability across sessions is still meaningful for explaining cognitive properties at the individual level. Third, to demonstrate applicability in a clinical sample, we investigated fingerprints and the correlation between fingerprint stability and intelligence in two samples with individuals with a psychotic disorder and healthy controls (OpenNeuro.ds000115 sample (Repovš and Barch, 2012) and the HCP Early Psychosis sample). Finally, to address the question of neural origins of the BOLD signal (Arthurs and Boniface, 2002), using data from the enhanced Nathan Kline Institute (eNKI) sample, we compared fingerprints between rs-fMRI and breath-holding scans that are known to capture cerebrovascular signals (Pinto et al., 2020). Altogether, our findings suggest that measures of local neural activity, and especially ReHo, show high specificity, sensitivity, and robustness as a neural fingerprint.

## Results

Analyses were performed using data from 399 unrelated subjects with two resting-state and seven task-based scans (acquired within 2 days) from the S1200 release of the Human Connectome Project Young Adult (HCP YA) sample (Feinberg et al., 2010; Moeller et al., 2010; Setsompop et al., 2012; Van Essen et al., 2013; Xu et al., 2021). The brain was parcellated into 264 spherical regions with a 3-mm radius around the Power coordinates (Power et al., 2011) (as analyses showed that smaller regions of interest (ROIs) yielded better identification accuracies - Supplementary Fig. S1). The global FC fingerprint was calculated as the Pearson correlations <u>between</u> the mean time series of each pair of ROIs (Finn et al., 2015) (Fig. 1). Furthermore, three local fingerprints were calculated: ReHo (Zang et al., 2004), ALFF (Zang et al., 2007), and fALFF (Zou et al., 2008). Distinct aspects of intrinsic brain function are captured by each of these measures: the often-reported FC fingerprint reflects the similarity between the mean time series of each pair of spatially distant ROIs (i.e. interregional similarity), while ReHo reflects the synchronization of the voxels’ time series <u>within</u> a ROI (i.e. intraregional similarity), and (f)ALFF reflects BOLD-fluctuations <u>within</u> a ROI (i.e. amplitude of intraregional activity). Next, identification accuracies, i.e., the fraction of correctly identified subjects, were calculated across all pairs of sessions. We observed near-perfect identification accuracies when using ReHo (mean: 99.6%), followed by ALFF (96%), fALFF (93%), and FC (84%) (Figure 2A), suggesting low intraindividual variability and high individual specificity. Next, we investigated whether our findings hold for different ways of performing identification.

**Figure 1.**
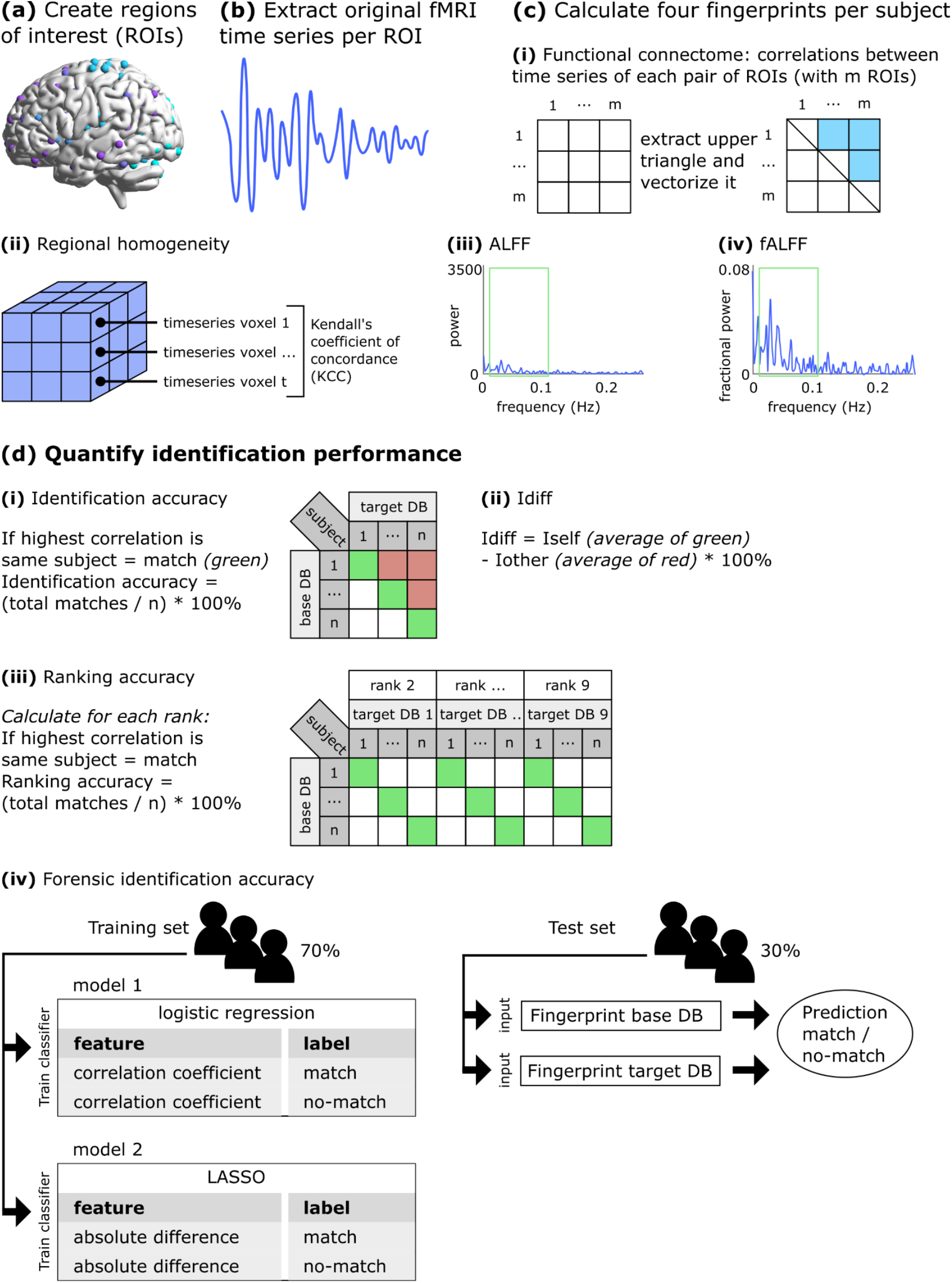
Schematic representation of identification methods.

**Figure 2.**
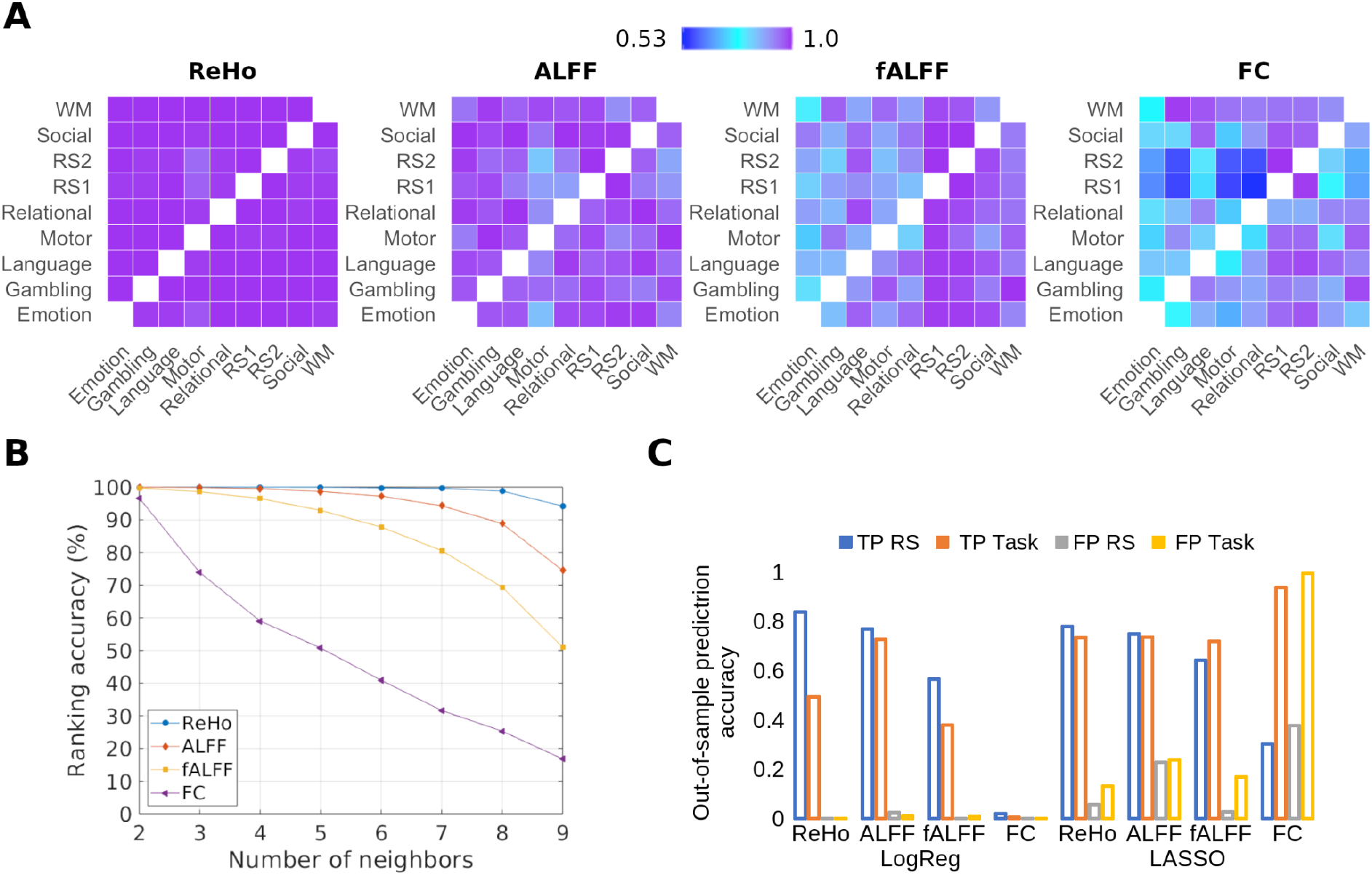
**A)** Comparison of identification accuracies of fingerprints in the HCP sample. **B)** Comparison of identification accuracies based on the ranking of fingerprints in the HCP sample. Note that at rank 2, the ranking accuracy is similar to the conventional identification accuracy, where the FC fingerprint performs relatively similarly to the local fingerprints, but the difference between them becomes evident at higher ranks. **C)** Forensic identification performance using fingerprint dissimilarity as a feature (LogReg) or the pattern of distances between the fingerprint elements as features (LASSO). For the FC, the nearly zero true-positive and false-positive rates with feature space 1 (fingerprint dissimilarity; left of Fig. 2C) indicate that it failed to match any of the test subjects, while the high true-positive and falsepositive rates with feature space 2 (pattern of distances between fingerprint elements; right of Fig. 2C) indicate that it matched many fingerprints correctly (true positives) but also incorrectly (false positives). Abbreviations: ReHo = regional homogeneity; ALFF = amplitude of low frequency fluctuations; fALFF = fractional ALFF; FC = functional connectome; RS = resting state; LogReg = logistic regression; TP = true positive rate; FP = false positive rate; HCP = Human Connectome Project.

### Highly accurate individual identification based on local functional fingerprints irrespective of identification performance quantification

First, we quantified the identification performance using the often-reported differential identifiability (“Idiff”) score. This score is the difference between the average within-subject and average between-subject similarity of a fingerprint; a higher Idiff indicates a better fingerprint (Amico and Goñi, 2018). Idiff scores were consistent with the accuracy results; the highest Idiff for ReHo (mean: 53), followed by ALFF (47), fALFF (38), and FC (10) (Supplementary Fig. S2; for general interest, additional results using a second parcellation (Dosenbach et al., 2010) are also reported). For those two conventional quantifications of identification performance, it is assumed that there is one fingerprint of a subject in the base database and one in the target database, and only the most similar fingerprint is considered a match. Here, we explored two alternative scenarios in which either multiple fingerprints or no fingerprint from the same subject exists in the target database. The first scenario was approached with a new score that we call *“ranking accuracy’,* which quantifies the degree to which fingerprints of *all sessions* of a subject are consistently ranked higher (i.e., show higher identification accuracy) than fingerprints of other subjects. This stricter measure yielded the same pattern of results (Figure 2B). For the second scenario in which there is the possibility of a “no-match”, we calculated the *“forensic identification’’* accuracy. The results clearly demonstrate the superiority of the local fingerprints (Fig. 2C).

### Better identification performance of local functional fingerprints is replicable across parcellations and samples and resilient to confounding effects

Additionally, we performed extensive evaluations to establish replicability and specificity of our results obtained with the Power 3-mm ROIs. First, we tested three other parcellations, namely, Dosenbach (Dosenbach et al., 2010), Shen (Shen et al., 2013), and Schaefer with 300 ROIs (Schaefer et al., 2018), which also showed higher identification accuracies for the local fingerprints (Fig. S3). Similar to our results with the Power spheres, the Dosenbach parcellation showed the highest identification accuracies for ReHo (94%), followed by ALFF (92%), fALFF (79%), and FC (78%). With the Shen and Schaefer parcellations, the results followed a similar pattern with a slight difference in the order: the highest identification accuracies were found for ALFF (77% and 79%), followed by ReHo (68% and 62%), FC (52% and 53%), and fALFF (37% and 48%). Second, we investigated the effect of removing confounds, i.e., sex, age, and total intracranial volume, from the fingerprints. Removal of these confounds increased overall accuracy while retaining the result pattern - the highest accuracy with ReHo (Fig. S4). Last, replication on the smaller Finn et al. (2015) HCP sample (n=125) showed similar results with some minor differences (see Supplementary Materials for details; Fig. S5 & S6).

### The dorsal attention and frontoparietal networks contribute most to individual uniqueness

Some brain networks contribute more to the “uniqueness” of the FC fingerprint than others (Finn et al., 2015; Jo et al., 2020). Encouraged by the high accuracy of the ReHo fingerprint, we investigated the contribution of each ROI in making the ReHo fingerprint unique. Each region was assigned to one of the seven Yeo networks based on the mode of the network membership of the voxels within that ROI (Yeo et al., 2011). Given that identification accuracies were highest between the resting state sessions but lowest with the motor task (Finn et al., 2015), data from these three sessions were used. We found that the ROIs contributing most to uniqueness were located in the dorsal attention network while the limbic network contained the fewest individual-specific ROIs (Fig. 3).

**Figure 3.**
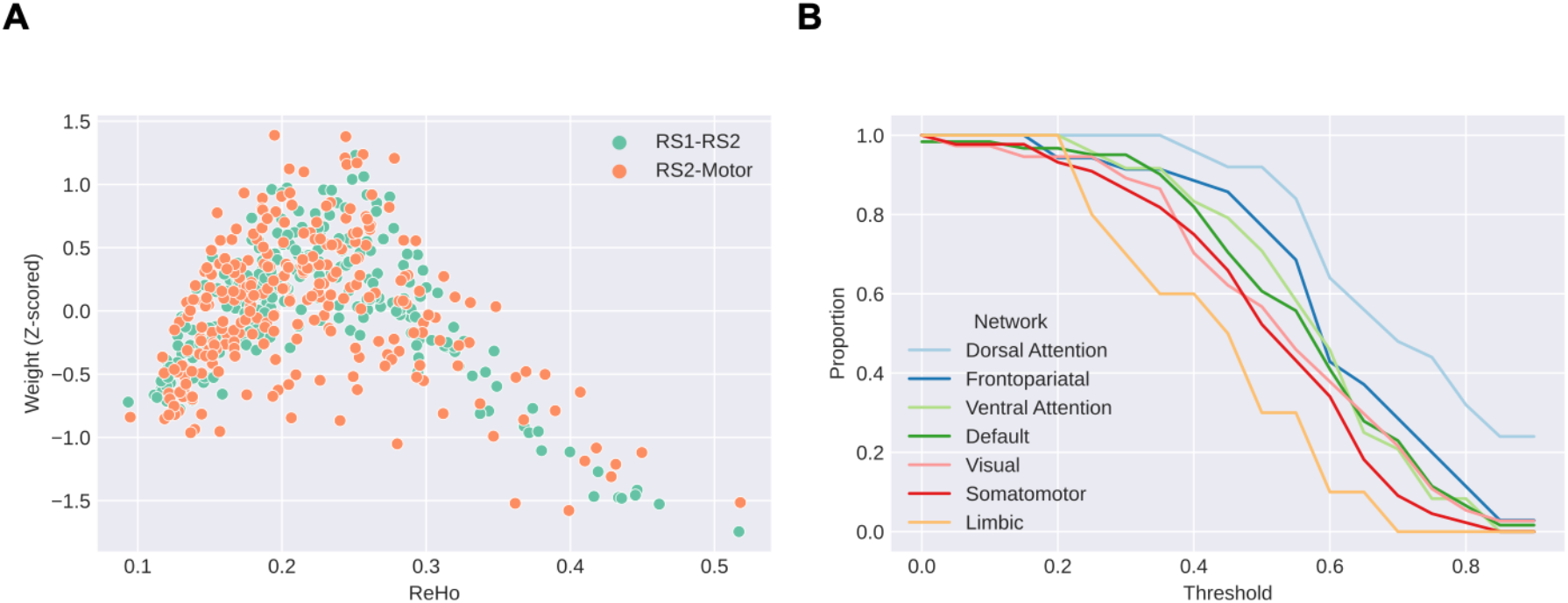
Contribution of regions in making the ReHo fingerprint individual-specific. **A)** Relationship between ReHo and Z-scored importance weights of regions averaged across all optimization runs. The average ReHo across subjects shows an inverse U-shaped relationship with the weights, with higher weights indicating higher individual uniqueness. **B)** The proportion of regions in Yeo’s 7 networks. Abbreviations: ReHo = regional homogeneity; RS1 = resting-state fMRI session day 1; RS2 = resting-state fMRI session day 2; Motor = fMRI motor task.

### Fingerprints and fingerprint stability capture interindividual differences

Next, for real-world and especially clinical applications, representations of brain function should not only have good fingerprint properties but should also capture interindividual differences in behavior or cognition. Interestingly, earlier studies have shown that the variation in the FC fingerprint measured across fMRI sessions, thus reflective of (task-related) reconfiguration, is lower in individuals with higher intelligence (Schultz and Cole, 2016; Thiele et al., 2021). Therefore, we chose to predict intelligence. First, we investigated the use of fingerprints to predict intelligence in the HCP dataset. We report findings if the median Pearson correlation between predicted and actual intelligence is higher than 0.15. We found that the ReHo fingerprint was predictive of crystallized (RS2 session, Social Cognition task) and fluid intelligence (Language task), the ALFF fingerprint was predictive of crystallized (Language task) and fluid intelligence (RS2 session, Motor task), the fALFF fingerprint of crystallized (Motor task) and fluid intelligence (Motor task), while the FC fingerprint was not predictive (Fig. S7-S15). Next, in addition to previous studies using only the fingerprint itself to predict cognition, we also examined the use of *the stability of the fingerprints* for the prediction of intelligence. This allowed us to test whether the (small) intraindividual fingerprint variability across sessions is still meaningful for explaining cognitive properties at the individual level. We observed that the stability of ReHo was predictive of crystallized intelligence, and ALFF stability predicted both crystallized and fluid intelligence.

In contrast, the stability of fALFF and FC was generally not predictive (Fig. 4A). A closer inspection of the LASSO weights for the ReHo features for predicting crystallized intelligence revealed that in individuals with higher crystallized intelligence the ReHo fingerprint was more stable between the resting state and the other sessions but less so between the language task and the other sessions. In addition, for the prediction of crystallized intelligence, high positive coefficients were observed for ALFF stability of the motor, language, and gambling tasks. For fluid intelligence, the motor and relational processing task’s ALFF stability received a high positive coefficient, and the social cognition task stability a negative coefficient.

**Figure 4.**
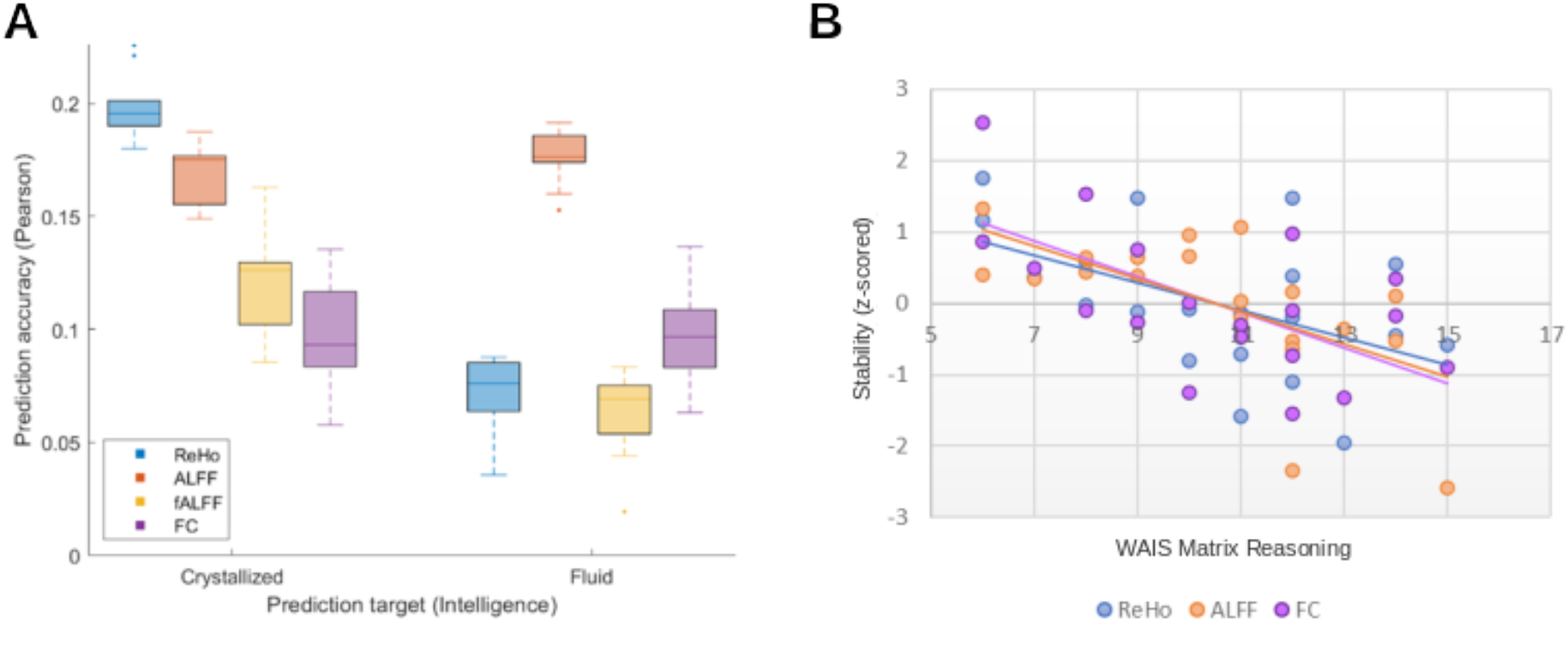
**A)** Prediction of crystallized and fluid intelligence using the stability of fingerprints of the nine sessions of the Human Connectome Project sample. Each boxplot shows the prediction performance across five cross-validation runs. **B)** Scatterplot between the WAIS Matrix Reasoning scores and the stability of the task fingerprints in the SZ sample of the OpenNeuro.ds000115 dataset. Abbreviations: ReHo = regional homogeneity; ALFF = amplitude of low frequency fluctuations; fALFF = fractional ALFF; FC = functional connectome.

### Local fingerprints are also stable in a clinical sample

In addition to the healthy population, neural fingerprints also need to be stable within clinical samples if they are to be used as clinical biomarkers. Studies have shown local abnormalities in schizophrenia (Shukla et al., 2010; Wang et al., 2019; Zhao et al., 2019), but it is unclear whether these abnormalities are stable. Kaufman et al. (2018) previously showed decreased FC fingerprint stability in patients with schizophrenia by combining three datasets (Kaufmann et al., 2018). Here, we used the publicly available OpenNeuro.ds000115 (Repovš and Barch, 2012) part of this previously reported sample, which includes 19 individuals with schizophrenia (SZ) and 20 healthy controls (see details in Supplementary Materials). fMRI data from the 0- and 2-back tasks obtained on the same day were used. This dataset does not contain rs-fMRI data. ROIs were again created with 3-mm radius spheres around the Power coordinates (Power et al., 2011). Identification accuracies for ReHo and ALFF were 100% for both controls and patients (analyzed separately), while they were lower in SZ for fALFF (65% in SZ; 88% in HC) and FC (99% in SZ; 100% in HC).

As this dataset does not contain rs-fMRI data, we additionally investigated the publicly available HCP Early Psychosis dataset. This dataset includes two resting-state fMRI sessions of 70 individuals with non-affective psychosis and 53 healthy controls (see details in Supplementary Materials). Identification accuracies were 100% for all metrics and both patients and HC (analyzed separately). This suggests that the ReHo and ALFF fingerprints are also stable in patients with psychotic disorders.

Next, we investigated the association between the stability of the fingerprints and intelligence in the schizophrenia sample and their matched healthy controls. Machine learning analysis, nor investigations of the association between fingerprints themselves and intelligence, were not attempted here due to the small sample size. Stability was calculated as the Pearson correlation between the fingerprints of the 0-back and 2-back task sessions (ds000115 sample) or resting state 1 and 2 sessions (HCP EP sample). In the ds000115 sample, we correlated the stability of the fingerprints with the Wechsler Adult Intelligence Scale (WAIS) Matrix Reasoning (variable WAIS_MATRICS_SCALE) and Vocabulary (variable WAIS_VOCAB_SCALED) subtest scores in both the SZ and HC groups. In the SZ group, the stability of the ReHo, ALFF, and FC fingerprints showed a high negative correlation with the WAIS Matrix Reasoning subtest (all r < −0.50, Fig. 4B), while in the HC group, only ALFF stability showed a positive correlation with WAIS Vocabulary subtest scores (r = 0.48) (Table S2). In the HCP EP sample, we correlated the stability of the fingerprints with the Fluid Cognition Composite Score (variable nih_fluidcogcomp_unadjusted) and the Crystallized Cognition Composite Score (variable nih_crycogcomp_unadjusted) while controlling for age. No significant correlations were found (see supplementary Table S3).

### ReHo represents a neural fingerprint

Altogether, thus far, our findings suggest that ReHo is most stable within an individual and captures brain-behavior associations. However, its high individual specificity might be driven by stable nonneural signals (i.e., artifacts, breathing, cardiovascular effects). Indeed, earlier studies have shown high stability of motion and cardiac pulsatility and that test-retest reliability of individual edges within the FC is reduced after artifact removal (Noble et al., 2019). Therefore, as the last follow-up analysis, we investigated the contribution of vascular activity to local and global fingerprints. Previous research showed that breath-holding is a reliable estimate of the influence of vascular activity on BOLD signals (Yuan et al., 2013), as it induces hypercapnia: an increase in carbon dioxide causes increased cerebral blood flow. This was investigated with the resting-state and breath-holding fMRI sessions (TR of 1400 ms) provided in the enhanced Nathan Kline Institute dataset (eNKI; n=478; see Supplementary Materials for details) (Nooner et al., 2012), as the HCP does not provide breath-holding fMRI data. The breath-holding paradigm consists of alternating breathing in, out, and breath-holding, causing a global change in oxygenation. We only used the breathholding blocks to derive the fingerprints. We observed the highest identification accuracies between resting state and breath-holding with ALFF (99%), followed by ReHo (87%), FC (72%), and fALFF (44%). Identification accuracies for ALFF and ReHo both reduced slightly when regressing out variance explained by ReHo and ALFF, respectively (ALFF: from 99% to 98%; ReHo: from 87% to 86%). This result suggests that the vascular signal in restingstate fMRI is captured with all fingerprints, but especially with ALFF.

## Discussion

Here, we compared the fingerprint properties of three local fingerprints with the commonly reported FC fingerprint, as higher reliability (lower intraindividual variability) of these local measures has been shown compared to the FC. Using a publicly available dataset and four different measures of identification performance, different parcellations, and ROI sizes, our findings show high accuracy of local functional fingerprints, especially when using ReHo. Next, four follow-up analyses were conducted. First, we show that the dorsal attention and frontoparietal networks contained the highest number of individual-specific ROIs in the ReHo fingerprint. Second, we demonstrate that intelligence can be predicted with local fingerprints and that even though the *intraindividual variability* of local fingerprints is small, it *is* meaningful given that it captures interindividual differences in intelligence. Third, using two other publicly available datasets we show the potential for the application of local fingerprints in clinical populations, as the ReHo fingerprint was highly accurate in individuals with a psychotic disorder, and its stability correlated with intelligence in individuals with schizophrenia. Finally, analyses using breath-holding data from yet another publicly available dataset suggest that the ReHo fingerprint largely reflects neuronal signals and that it can indeed be deemed a “neural fingerprint”. As such, our findings suggest the ReHo fingerprint as a good candidate for further exploration: it has excellent fingerprint properties and is predictive of interindividual differences in cognition.

### “Fingerprinting” with local fingerprints versus the functional connectome

ReHo quantifies the synchronization among the voxelwise time series within a ROI, and its stability across sessions appears to reflect the existence of regions that work relatively uniformly across mental states (i.e., tasks). The FC, on the contrary, captures dynamic and state-dependent interactions between regions and might be less suitable for answering identifiability questions. One other explanation for our finding of higher accuracy of the local fingerprints could be that first-order statistics such as ReHo and (f)ALFF might show different patterns of interindividual variation compared to second-order statistics such as the FC (Liégeois et al., 2017). Moreover, for local metrics, the values are averaged for a region, which reduces noise (Xu et al., 2016). To the best of our knowledge, no studies have reported identification accuracies for ReHo or (f)ALFF fingerprints thus far. Two studies reported identification accuracies for a different measure of local brain function called BOLD variance (Finn et al., 2015) or BOLD variability (BV) (Gaut et al., 2019). The BV fingerprint reflects the BOLD variance per region, i.e., regional fluctuations similar to the ALFF fingerprint but in the full frequency range. Both of these studies showed lower identification accuracies of the BV fingerprint compared to the FC fingerprint, suggesting that not *all* local fingerprints have better fingerprint properties than the FC and that the BV fingerprint captures processes differently from ALFF.

### The dorsal attention and frontoparietal networks contribute most to individual “uniqueness”

Previous studies predominantly showed the contribution of frontoparietal regions or “higher-order networks” (i.e., frontal, parietal, and temporal lobes) to FC identifiability (Airan et al., 2016; Amico and Goñi, 2018; Cai et al., 2021; Finn et al., 2015; Miranda-Dominguez et al., 2014). For example, one study found that ROIs contributing most to the “uniqueness” of the FC were located in the default mode, attention, and executive control networks (Airan et al., 2016). Here, we investigated “uniqueness” for the ReHo fingerprint by optimally weighting ReHo features higher in some brain regions to enhance the fingerprint efficacy, thereby revealing brain regions that contribute more to individual uniqueness. Next, each region was assigned to one of the previously described seven networks based on the mode of the network membership of the voxels within that ROI (Yeo et al., 2011). We found that regions in the dorsal attention and frontoparietal networks contributed the most to identifiability, while the limbic network contributed the least. However, some critical considerations should be noted. First, limbic regions in these networks are smaller than ROIs within the other networks and therefore might be affected more by normalization errors that induce noise. Second, they are also affected most by magnetic susceptibility artifacts causing a reduced signal-to-noise ratio leading to reduced homogeneity (Aiello et al., 2015), which can make the ReHo fingerprint noisier and hence less specific in these regions. Last, the nature of the fMRI sessions used to investigate uniqueness (the motor task and resting state sessions) might also influence the contribution of certain ROIs to the uniqueness of the fingerprint (Barch et al., 2013).

Previous research has indeed shown greater interindividual variability of multimodal association networks (i.e., default, dorsal attention, and executive control) (Laumann et al., 2015; Mueller et al., 2013; Xu et al., 2016), while unimodal networks (i.e., visual and sensorimotor) had lower interindividual variability (Xu et al., 2016). On the other hand, the DMN, attention, and subcortical systems are relatively less dynamically diverse (Bansal et al., 2019) and thus show lower intraindividual variability, while unimodal networks have higher intraindividual variability (Xu et al., 2016). It appears that higher-order networks are affected more by genetic and environmental factors (Gao et al., 2014; Xu et al., 2016), while lower-order networks are influenced more by situational or task demands (Xu et al., 2016). Altogether, our findings suggest that not only the *distributed connectivity* but also the *local brain function* within higher-order brain networks contributes most to the “uniqueness” of a brain. These networks that are stable within an individual but highly variable across individuals might provide a fundamental backbone of functional brain organization implicated in cognition as well as in disease (Bansal et al., 2019). Future studies might use the localizing power of local fingerprints to attempt to pinpoint “uniqueness” to even smaller regions.

### Fingerprints and fingerprint stability capture interindividual differences

Our findings thus far showed that the ReHo fingerprint is an accurate neural fingerprint. Next, we showed that the local fingerprints are predictive of crystallized and fluid intelligence, while the FC fingerprint is not predictive. Previous studies significantly predicted fluid intelligence from the FC fingerprint itself (Finn et al., 2015) and the strength of the dynamic FC fingerprint (Liu et al., 2018), a high-resolution FC fingerprint (Mansour L et al., 2021), and a FC fingerprint refined using an autoencoder network (Cai et al., 2021). These studies did not attempt to predict crystallized intelligence (i.e., the composite score of the Picture Vocabulary Test and Oral Reading Recognition Test). Only the last-mentioned study investigated one crystallized intelligence subtest, the Picture Vocabulary Test - measuring language comprehension - and found that the refined FCs could significantly predict it (Cai et al., 2021). The predictiveness of local fingerprints has not been investigated, although a few studies found significant associations between (voxelwise) whole-brain ReHo or ALFF and intelligence (Li et al., 2018; Wang et al., 2011). In short, our findings suggest that different fingerprints will be more appropriate for predicting different traits, just as some fMRI tasks are more appropriate for capturing interindividual differences in specific traits (Greene et al., 2018).

The fingerprints based on the Motor fMRI task data appeared to be most predictive of intelligence, which is also the session with the lowest identification accuracies. Indeed, it has been shown that reliability is not correlated with predictiveness (Noble et al., 2017) (of FC edges in the referenced study), conceivably because behavior is not static but dynamic (Finn and Rosenberg, 2021). Therefore, we next tested whether the subtle differences in the fingerprints across sessions are still informative of individual-level intelligence. Using a machine learning framework, we predicted intelligence with the stability of the fingerprints across sessions. Our results showed that the stability of ReHo could predict crystallized intelligence but not fluid intelligence. On the other hand, ALFF stability could predict both crystallized and fluid intelligence, while the stabilities of fALFF and FC fingerprints were not predictive. Thus, even though local fingerprints are highly similar across sessions, the (small) variation due to changes in mental state is still predictive of individual-level cognition.

Our findings suggest that intraregional information processing, as captured with ReHo, is stable across tasks or mental states. This stability of information processing *within* a region is associated more with the more static form of intelligence that is conceptualized with crystallized intelligence than with the more dynamic ability to reason and solve new problems referred to as fluid intelligence, which might be associated more with brain dynamics or interactions *between* regions. In a previous study in which functional connectivity patterns were also more predictive of crystallized than fluid intelligence, the authors suggested that this might be explained by the strong mapping between brain anatomy and language, while fluid intelligence might rely on the functioning of multiple (partly) overlapping brain networks that are involved in many distinct cognitive skills (Dhamala et al., 2021). Our findings show that ALFF was predictive of both crystallized and fluid intelligence. ALFF reflects temporal variability and therefore might be more suited for capturing dynamic changes or behaviors. Along this line, a study examining time-varying network configurations found that dynamic configurations are not only good fingerprints in healthy individuals but that the individual variability in one of the configurations (i.e., brain states) was predictive of florid psychotic symptoms in individuals with a psychotic disorder (Reinen et al., 2018). Future studies could systematically investigate the ability of different fingerprints to predict different types of behavior and cognitive functions.

A closer inspection of the LASSO weights for the ReHo features for predicting crystallized intelligence revealed that reconfigurations of ReHo across states were smaller in individuals with higher crystallized intelligence. This is in line with studies showing that the FC fingerprint at rest was more similar to the FC during several tasks in individuals with higher intelligence (Schultz and Cole, 2016; Thiele et al., 2021). This ability to reconfigure networks efficiently based on task demands can be seen as a sign of high intelligence (Schultz and Cole, 2016). Surprisingly, we also found that *higher* reconfiguration of the ReHo fingerprint between the language task and the other sessions was predictive of higher crystallized intelligence. However, some differences can be noted compared to this previous study using the HCP dataset, which may explain the difference in findings: they examined the FC fingerprint and not the ReHo fingerprint, calculated stability between fingerprints with the cosine distance between FC matrices, and calculated intelligence with a latent g-factor derived from 12 cognitive scores (Thiele et al., 2021).

### Local fingerprints are also stable in a clinical sample

The application of biomarkers in precision psychiatry requires stability in clinical populations. Here, using the OpenNeuro.ds000115 dataset, we show perfect identification accuracies (100%) for ReHo and ALFF in both SZ and HC. For fALFF and FC, identification accuracies for SZ were lower compared to HC, albeit very slightly for FC. The higher identification accuracies for the FC in this sample compared to the HCP YA sample (99-100% vs 54-100%) could be due to the smaller sample size (19 SZ and 20 HC vs 399 HC), longer task duration compared to some (but not all) HCP sessions (+/− 4 minutes vs +/− 2-14,5 minutes), and the (increased) engagement of similar networks during 2- and 0-back tasks, all of which have been shown to increase FC fingerprint identification accuracy. With the HCP Early Psychosis dataset, we show perfect identification accuracies (100%) for all fingerprints in both patients with non-affective psychosis and HC. This is in line with our findings in the HCP YA sample in which identification accuracies using two resting state sessions were also perfect for all fingerprints.

The utility of FC fingerprints has been shown in several clinical populations. A delay in reaching FC fingerprint stability has been reported even in individuals with preclinical signs of mental illness (i.e., increased prodromal symptoms across domains of ADHD, schizophrenia, and depression) (Kaufmann et al., 2017). Moreover, some aspects of FC functioning are disrupted - *and* associated with symptom severity - *across* patients with different psychiatric diagnoses, such as primary psychotic disorder, affective psychosis, and primary affective disorder without psychosis, while other aspects were only disrupted in patients with a psychotic disorder specifically (Baker et al., 2019). Lower stability of the whole-brain FC (between 0- and 2-back fMRI tasks) has also been shown in schizophrenia patients (Kaufmann et al., 2018). Altogether, these previous studies demonstrated the transdiagnostic utility of FC fingerprints. While this was unknown for local fingerprints, our findings suggest the utility of local functional fingerprints in individuals with psychotic disorders. Future studies taking a transdiagnostic approach could inform the stability of different functional fingerprints across individuals with different characteristics and their ability to predict different types of behaviors and symptoms.

Next, we investigated the association between fingerprint stability and intelligence. Given the small sample size, machine learning analysis to examine the fingerprints’ ability to *predict* intelligence was not attempted here. We found that in individuals diagnosed with schizophrenia from the OpenNeuro.ds000115 dataset, *higher* reconfigurations of ReHo, ALFF, and FC fingerprints were associated with higher scores on the Matrix Reasoning subtest, a measure of abstract problem-solving and inductive reasoning, which is considered a part of fluid intelligence. This finding was opposite to what we observed in healthy individuals of the HCP YA sample, which was also not seen in the matched HC of the OpenNeuro.ds000115 dataset (all r < 0.02-0.1). This finding could suggest that individuals with SZ had difficulties performing the 2-back task and failed to engage the networks required for the additional working memory load of this task (compared to the 0-back task). Indeed, a previous study using the same dataset (including 19 SZ and 10 HC) showed lower 2-back performance in SZ compared to HC (0.72 vs 0.81 for 0-back; 0.64 and 0.80 for 2-back) (Repovš and Barch, 2012). In HC, only ALFF stability was associated with the scores on the Vocabulary subtest. In contrast to the OpenNeuro.ds000115 dataset, with the HCP Early Psychosis dataset, no significant correlations were found between fingerprint stability and intelligence. This might be explained by several factors that differ between the two datasets: 1) the HCP EP sample consists of individuals in an early phase of psychosis while the OpenNeuro.ds000115 sample consists of patients with longer illness duration, 2) intelligence was measured differently, and 3) use of resting-state fMRI sessions with a lower variation of fingerprint stability in the HCP EP dataset in contrast to task fMRI sessions in the OpenNeuro.ds000115 dataset (see Fig. S18). Altogether, our findings suggest that even though the ReHo and ALFF fingerprints are highly stable, their subtle variation due to mental states is informative of individual-level cognition in patients with schizophrenia.

### ReHo represents a neural fingerprint

Higher ReHo, fALFF (Aiello et al., 2015), and ALFF (Tomasi et al., 2013) have been associated with higher glucose metabolism, although the strength of this association varies across the brain. As the fourth follow-up analysis, we investigated the contribution of vascular activity to local and global fingerprints. Identification accuracies with the eNKI breath-holding fMRI-task and resting-state data suggest that breathing, or vascular signal, is represented more in resting-state ALFF than in resting-state ReHo. Indeed, previous research has shown that ALFF captures this signal (Golestani et al., 2016; Zuo et al., 2010). Furthermore, our results are in line with the idea that fALFF, as a normalized ALFF, is affected less by physiological signals from large blood vessels and CSF that affect the entire frequency spectrum. Taken together, the high accuracy of ReHo in the HCP samples and its relatively lower accuracy with breath-holding data suggests that, as expected, ReHo captures neural signals more than ALFF.

### Better identification performance of local functional fingerprints is replicable across parcellations and samples and resilient to confounding effects

Finally, we tested the robustness of our main results regardless of methodological choices. In addition to the results obtained with ROIs created with 3-mm spheres around the Power coordinates, identification was also evaluated using three other parcellations (i.e., Dosenbach, Shen, and Schaefer with 300 ROIs). While the results with the Dosenbach spheres were similar to the Power spheres, for the Shen and Schaefer parcellations, higher identification accuracies were found for ALFF, followed by ReHo, FC, and fALFF (versus ReHo, ALFF, fALFF, FC with Power and Dosenbach parcellations). This difference might be explained by the fact that ROIs in the Power (264 ROIs; 3-mm radius) and Dosenbach (160 ROIs; 5-mm radius) parcellations are compact spheres, while the ROIs of the Shen (268 ROIs) and Schaefer (300 ROIs) parcellations are much larger. As we calculated ReHo for each ROI as Kendall’s coefficient of concordance for all the voxelwise time series within that ROI, ReHo values are expected to converge to 0 with increasing ROI size, in effect rendering the ReHo fingerprint less specific and confirming the importance of spatial resolution in ReHo calculation. fALFF values increase with larger ROI sizes, as larger ROIs result in smoother signals and therefore a higher proportion of low-frequency fluctuations (Zhang et al., 2021). Moreover, larger ROIs reflect more macroscopic organization, and are impacted more by partial volume effects, and averaging within them deteriorates local fingerprints. While a previous study showed lower identification accuracies with larger ROI sizes for the FC fingerprint (Li et al., 2021), our findings show that this also applies to local fingerprints. Next, we tested the effect of sample size, as larger sample sizes are essential for application in precision psychiatry, while it is expected that they yield lower identification accuracies (Li et al., 2021). Therefore, all comparisons were also conducted with the previously reported sample of 125 subjects (Finn et al., 2015). We found that identification accuracies for the FC decreased the most with increasing sample size, while local fingerprints, especially ReHo, were affected the least. Last, our finding of highly accurate local fingerprints was resilient against confounding effects (i.e., age, sex, and intracranial volume), as confound removal resulted in the same pattern of identification accuracies, with ReHo yielding the highest accuracies. Confound removal benefited identification accuracies for FC the most and ALFF the least, suggesting higher sensitivity of the FC and lower sensitivity of ALFF to confounding effects. Altogether, our findings suggest that ReHo shows high specificity, sensitivity, and robustness as a neural fingerprint.

### Limitations

This study has some limitations. First, the contribution of physiological signals to the BOLD signal is unclear. We tested the contribution of vascular activity to the fingerprints, but other sources of physiological signals might also contribute. However, we used the FIX-denoised resting-state data from the HCP, which is expected to be free of such extraneous signals. Moreover, several studies have shown that accurate subject identification is mostly based on neural signals and not on nuisance processes such as physiological processes and head motion (Airan et al., 2016; Xifra-Porxas et al., 2021). Second, the contribution of structural information (brain morphology) to fingerprint identification accuracy is unclear (Gaut et al., 2019). Third, we created fingerprints based on local brain function, which is just one of many possibilities. Other studies investigated fingerprints based on shape, white matter fiber geometry, voxelwise diffusion density, or combinations (Kumar et al., 2018). Future studies could also investigate a multimodal fingerprint including ReHo. Fourth, we analyzed data that was acquired within a short period (i.e., days), but previous research has shown decreasing identification accuracies with increasing periods (i.e., from 90% within one month to 66% within 2-3 years) (Ni et al., 2021). Future work could examine the stability of local and global fingerprints over months or years using longitudinal data. Fifth, we did not control for differences in scan durations between rs- and task sessions. However, earlier research has shown that identification accuracies between task sessions improved when matching scan duration but were still highly variable across sessions (Finn et al., 2017). Sixth, we checked fingerprint stability in two clinical samples of individuals with psychotic disorders. It is unclear whether our findings generalize to other clinical samples and conditions. Finally, local metrics such as ReHo and (f)ALFF were originally proposed for rs-fMRI data while we also calculated them with task-based fMRI data. However, this data was treated as resting state, as task timing was ignored. Our findings show that ReHo and (f)ALFF are highly stable across resting state and tasks, suggesting that even when influenced by a task they appear to reflect intrinsic brain function.

### Conclusion

Investigating associations between brain function and cognition, behavior, or disease characteristics requires within-individual stable representations of brain function (Tipnis et al., 2020). Recent studies have increasingly stressed the importance of neural representations capturing both intraindividual stability and interindividual differences, as both are desired properties for clinical biomarkers (Finn and Rosenberg, 2021; Mantwill et al., 2021). Here, we show that measures of local brain function - and especially ReHo - are better fingerprints compared to the global FC fingerprint, while they also capture interindividual differences in cognition. We suggest the ReHo fingerprint to be a good candidate for further exploration of applicability in precision medicine.

## Materials and Methods

### Calculation of identification accuracy in the Human Connectome Project sample

#### Human Connectome Project (HCP) sample

Data collection was approved by the Institutional Review Board at Washington University in St. Louis. The selection of subjects was performed in R version 3.6.1 (R Core Team 2019). Individuals were included in case of the availability of 1) a T1-weighted scan, 2) two restingstate fMRI scans (rfMRI_REST1 and rfMRI_REST2), 3) data on 58 behavioral variables (Kong et al., 2019) and 4) all fMRI tasks (working memory (tfMRI_WM), gambling (tfMRI_GAMBLING), motor (tfMRI_MOTOR), language (tfMRI_LANGUAGE), social cognition (tfMRI_SOCIAL), relational processing (tfMRI_RELATIONAL) and emotion processing (tfMRI_EMOTION) (Barch et al., 2013). Subjects with reported issues according to the HCP data release updates were excluded. From this sample, subjects were selected that were unrelated (variable Family_ID) with a similar number of selected males and females (Gender) that were matched on age (Age_in_Yrs), education (SSAGA_Educ), race (Race), and BMI (BMI). This resulted in a sample of 399 subjects consisting of 195 females and 204 males. The mean age, education, and BMI were similar between males and females, namely, 27.83 (sd=3.75) versus 29.68 (sd=3.69), 14.08 (sd=1.75) vs 15.01 (sd=1.86), and 27.03 (sd=4.38) vs 26.06 (sd=5.59), respectively. The distribution of races was also similar between males and females: Asian/Nat. Hawaiian/Other Pacific Is. (11 vs 13), Black or African Am. (28 vs 29), White (154 vs 145), unknown or not reported (4 vs 2), more than one (7 vs 6). For all subjects, all available fMRI data (i.e., two resting-state fMRI sessions and seven fMRI tasks) were used for the main analyses. Further details on the tasks and acquisition parameters can be found in Barch et al., 2013 (Barch et al., 2013).

We used the preprocessed data provided by the HCP, which includes artifact removal, motion correction, and registration to standard space (Fischl, 2012; Glasser et al., 2013; Jenkinson et al., 2012, 2002; Woolrich et al., 2001). For the resting-state data, the ICA-FIX noise components removed data were used. We also used the sample from Finn et al. (Finn et al., 2015) (n=125) for replication and optimization analyses; of this sample, 47 subjects overlapped with the n=399 HCP subsample.

#### Preprocessing

Additional minimal preprocessing was performed assuming no prior knowledge of the data, therefore allowing the use of our preprocessing pipeline in a broad range of situations (i.e., task- or resting-state data, variable scan parameters, etc.). This was done in MATLAB release 2019b (MathWorks) and included regressing out the 12 detrended motion parameters (provided by HCP as Movement_Regressors_dt.txt), as well as the mean time courses of WM, CSF, and global signal, and filtering the time series with a bandpass filter of 0.01-0.1 Hz. Next, the fingerprints were calculated for both left-right (LR) and right-left (RL) phase-encoding runs and consequently averaged.

#### Local fingerprints

A local fingerprint for a subject consists of a vector of length N, where N is the number of ROIs. All local fingerprints were calculated on the average time series per ROI. ReHo is calculated with Kendall’s coefficient of concordance considering the time series of all the voxels within an ROI (Zang et al., 2004). For ALFF and (f)ALFF, voxelwise time series were transformed to the frequency domain to obtain the power spectrum (Zou et al., 2008). ALFF is the average square root of the power in the 0.01-0.1 Hz frequency range averaged across voxels within a ROI (Zang et al., 2007). fALFF is ALFF scaled by the total power in the full frequency range averaged across voxels within a ROI (Zou et al., 2008).

#### Global fingerprint (whole-brain functional connectome)

A FC for a subject consists of a matrix of size NxN, where N is the number of ROIs. We extracted the lower triangle of this matrix and vectorized it to obtain the FC fingerprint as a vector of length N x (N-1)/2.

#### Identification accuracy

Iteratively, all subjects’ data from one session was used as the “base dataset” while data from another session was used as the “target dataset”. Spearman correlations were calculated between the fingerprints of the two sessions. Using a 1-nearest neighbor approach (Finn et al., 2015), the “base” fingerprint was labeled with the subject identity of the most similar fingerprint in the “target dataset”. Identification accuracy was calculated as the fraction of correctly labeled participants.

### Idiff

An identification matrix of size NxN (N = number of subjects) was created reflecting the correlations between the fingerprints of a subject in the base and all fingerprints in the target database. The lower triangle elements were extracted and represented as an N x (N-1)/2-dimensional vector. “Iself” is the average within-subject similarity, while “Iothers” is the average between-subjects similarity. Idiff is then calculated by taking the difference between Iself and Iothers (Amico and Goñi, 2018).

### Ranking accuracy

Identification accuracies were calculated (see paragraph “Identification accuracy” above) with one difference. Namely, iteratively, each session was used as the base dataset, while <u>all other eight sessions</u> (instead of only one other session) were included in the target dataset. Then, instead of focusing only on the fingerprint in the target dataset with the highest correlation, all fingerprints in the target dataset were ranked based on their similarity with a subject’s fingerprint in the base dataset. At each rank (2-9, as each subject has data from nine sessions), the proportion of correctly identified subjects was calculated (see Fig. 1 for a visualization).

### Forensic identification

We used a machine learning framework to predict whether two fingerprints matched (see Fig. 1). For this, we used the two resting-state sessions as training data and trained a classification model using a match-no-match dataset derived from 70% of the subjects (randomly chosen). Forensic identification accuracies were calculated across all pairs of sessions. We tested two different feature spaces that were constructed containing a match and a no-match part. The match part contained features of fingerprints from two sessions of the same subject. The no-match part contained features of a subject from session 1 and their second-closest match from session 2. For the first model, the feature space included (for each subject) the correlation coefficient between two resting-state fingerprints. For the second model, the features for the match-part included (for each subject) the absolute difference between the elements of two resting-state fingerprints. Similarly, the features for the no-match part included the absolute difference between the resting state 1 fingerprint of a subject in the base dataset and the resting state 2 fingerprint of the closest other subject in the target dataset. This provided a dataset with 2xN rows (N = number of subjects), which together with class labels indicating match or no-match was used to train a classifier. More specifically, we employed logistic regression with the feature space of the first model and LASSO with the feature space of the second model. The 30% hold-out subjects were used for testing: each of the 30% held-out subjects was probed against all the subjects one by one as a reference to obtain matches. Given a reference fingerprint (with known identity) and a test fingerprint (unknown identity), a trained model was used to predict whether they matched. A prediction was then obtained using each fingerprint in a reference dataset. The reference dataset was either the average of the two resting-state sessions or one of the task sessions (emulating the privacy-preserving scenario where the training data itself is not available). As the number of predictions for each test fingerprint equals the number of fingerprints in the reference dataset, predictions for a test fingerprint can contain one, multiple, or no matches. If the correct subject was predicted as a match, it was counted as a true positive (TP), and if any other subject was predicted as a match (irrespective of TP), then this was counted as a false positive (FP). A good classifier should provide a high TP and a low FP rate. The split between training and test subjects was repeated 10 times. Ten times repeated 10-fold cross-validation was performed on the training data to estimate the generalization ability.

### Confound removal

Confounding signals due to age, sex, and total intracranial volume (ICV) were regressed out from each column of each database (subjects x fingerprint), and the residuals were used for identification. The ICV was estimated using the CAT12.5 VBM processing pipeline.

### Contribution of regions to individual uniqueness via *margin optimization*

To identify important ROIs whose ReHo is individual-specific, we posed an optimization problem to derive weights for the weighted Pearson correlation coefficient that maximizes the margin, i.e., the correlation difference between the correct fingerprint and the next best. As the ReHo identification accuracy was very high (close to 100% in most cases), using the accuracy as an optimization objective would not be meaningful. We sought to identify a weight w_i_ for each ROI by solving the following optimization problem:

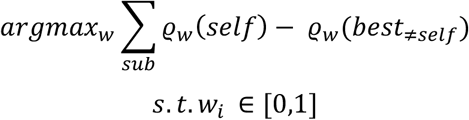

This optimization problem was solved to maximize the average margin for the HCP sample (n=399) using the CMA-ES algorithm (Hansen and Ostermeier, 2001). Importantly, the data was divided into a 50% training set used to learn the weights and a 50% test set to validate whether the optimized weights indeed improve the margin on out-of-sample data. This procedure was repeated 20 times to assess stability. We used two optimization problems: RS1-RS2 and RS2-Motor. This resulted in an importance weight for each ROI reflecting its contribution to uniqueness (Fig. S16 & S17). Next, each of the Power 3-mm ROIs was assigned to one of the seven Yeo networks or ‘None’ based on the mode of the network membership of the voxels within each ROI. Then, we varied a threshold over the optimized weights (averaged over RS1-RS2 and RS2-Motor) to assess the retained proportion of the ROIs from each of the seven networks (Table S1).

### Fingerprints and fingerprint stability predict intelligence

The fingerprints were used to predict individual-level crystallized (“CogCrystalComp_AgeAdj”) and fluid intelligence (“CogFluidComp_AgeAdj”) separately in a 5 times repeated 5-fold cross-validation setup using least absolute shrinkage and selection operator (LASSO) regression (Hastie et al., 2009). The variance due to age and total intracranial volume was removed from the features in a cross-validation consistent manner (More et al., 2021).

Next, to investigate the predictiveness of fingerprint *stability,* we first calculated - for each individual - the stability of the fingerprints as the average Pearson correlation of each session’s fingerprint with the fingerprints of all other sessions resulting in a feature vector with nine values (i.e., two resting-state and seven task fMRI sessions). These features were then employed for prediction analysis as described above.

## Acknowledgments

The authors would like to thank Dr. Branislava Ćurčić-Blake for helpful discussions and Dr. Emily S. Finn for valuable comments on a previous version of this manuscript. Data were provided [in part] by the Human Connectome Project, WU-Minn Consortium (Principal Investigators: David Van Essen and Kamil Ugurbil; 1U54MH091657) funded by the 16 NIH Institutes and Centers that support the NIH Blueprint for Neuroscience Research; and by the McDonnell Center for Systems Neuroscience at Washington University. Data and/or research tools used in the preparation of this manuscript were obtained from the National Institute of Mental Health (NIMH) Data Archive (NDA). NDA is a collaborative informatics system created by the National Institutes of Health to provide a national resource to support and accelerate research in mental health. Dataset identifier: http://dx.doi.org/10.15154/1528137. This manuscript reflects the views of the authors and may not reflect the opinions or views of the NIH or of the Submitters submitting original data to NDA.

## Funding

This work was supported by the Deutsche Forschungsgemeinschaft, The Helmholtz Portfolio Theme ‘Supercomputing and Modelling for the Human Brain’ and the European Union’s Horizon 2020 Research and Innovation Programme [grant number 945539 (HBP SGA3) and grant number 826421 (VirtualBrainCloud) to SBE].

## Data and materials availability

### Code availability

Code is available on GitHub (link will be added later).

### Data availability

Used datasets are publicly available: the HCP Young Adult dataset (https://www.humanconnectome.org/study/hcp-young-adult) and the HCP Early Psychosis dataset (https://www.humanconnectome.org/study/human-connectome-project-for-early-psychosis) require signing a data use agreement, eNKI (http://fcon_1000.projects.nitrc.org/indi/enhanced/), and OpenNeuro.ds000115 (https://openneuro.org/datasets/ds000115/versions/00001).

## Notes

### Competing Interest Statement

The authors have declared no competing interest.

### Summary of Updates

Analyses and results using the HCP Early Psychosis sample are added

## References

Aiello, M., Salvatore, E., Cachia, A., Pappatà, S., Cavaliere, C., Prinster, A., Nicolai, E., Salvatore, M., Baron, J.-C., Quarantelli, M., 2015. Relationship between simultaneously acquired resting-state regional cerebral glucose metabolism and functional MRI: a PET/MR hybrid scanner study. Neuroimage 113, 111–121. doi:10.1016/j.neuroimage.2015.03.017

Airan, R.D., Vogelstein, J.T., Pillai, J.J., Caffo, B., Pekar, J.J., Sair, H.I., 2016. Factors affecting characterization and localization of interindividual differences in functional connectivity using MRI. Hum. Brain Mapp. 37, 1986–1997. doi:10.1002/hbm.23150

Amico, E., Goñi, J., 2018. The quest for identifiability in human functional connectomes. Sci. Rep. 8, 8254. doi:10.1038/s41598-018-25089-1

Arthurs, O.J., Boniface, S., 2002. How well do we understand the neural origins of the fMRI BOLD signal? Trends Neurosci. 25, 27–31. doi:10.1016/s0166-2236(00)01995-0

Baker, J.T., Dillon, D.G., Patrick, L.M., Roffman, J.L., Brady, R.O., Pizzagalli, D.A., Öngür, D., Holmes, A.J., 2019. Functional connectomics of affective and psychotic pathology. Proc Natl Acad Sci USA 116, 9050–9059. doi:10.1073/pnas.1820780116

Bansal, K., Garcia, J.O., Tompson, S.H., Verstynen, T., Vettel, J.M., Muldoon, S.F., 2019. Cognitive chimera states in human brain networks. Sci. Adv. 5, eaau8535. doi:10.1126/sciadv.aau8535

Barch, D.M., Burgess, G.C., Harms, M.P., Petersen, S.E., Schlaggar, B.L., Corbetta, M., Glasser, M.F., Curtiss, S., Dixit, S., Feldt, C., Nolan, D., Bryant, E., Hartley, T., Footer, O., Bjork, J.M., Poldrack, R., Smith, S., Johansen-Berg, H., Snyder, A.Z., Van Essen, D.C., WU-Minn HCP Consortium, 2013. Function in the human connectome: task-fMRI and individual differences in behavior. Neuroimage 80, 169–189. doi:10.1016/j.neuroimage.2013.05.033

Cai, B., Zhang, G., Zhang, A., Xiao, L., Hu, W., Stephen, J.M., Wilson, T.W., Calhoun, V.D., Wang, Y.-P., 2021. Functional connectome fingerprinting: Identifying individuals and predicting cognitive functions via autoencoder. Hum. Brain Mapp. 42, 2691–2705. doi:10.1002/hbm.25394

Chen, B., Xu, T., Zhou, C., Wang, L., Yang, N., Wang, Z., Dong, H.-M., Yang, Z., Zang, Y.-F., Zuo, X.-N., Weng, X.-C., 2015. Individual Variability and Test-Retest Reliability Revealed by Ten Repeated Resting-State Brain Scans over One Month. PLoS ONE 10, e0144963. doi:10.1371/journal.pone.0144963

Chen, J., Tam, A., Kebets, V., Orban, C., Ooi, L.Q.R., Marek, S., Dosenbach, N., Eickhoff, S., Bzdok, D., Holmes, A.J., Thomas Yeo, B.T., 2020. Shared and unique brain network features predict cognition, personality and mental health in childhood. BioRxiv. doi:10.1101/2020.06.24.168724

Dhamala, E., Jamison, K.W., Jaywant, A., Dennis, S., Kuceyeski, A., 2021. Distinct functional and structural connections predict crystallised and fluid cognition in healthy adults. Hum. Brain Mapp. 42, 3102–3118. doi:10.1002/hbm.25420

Dosenbach, N.U.F., Nardos, B., Cohen, A.L., Fair, D.A., Power, J.D., Church, J.A., Nelson, S.M., Wig, G.S., Vogel, A.C., Lessov-Schlaggar, C.N., Barnes, K.A., Dubis, J.W., Feczko, E., Coalson, R.S., Pruett, J.R., Barch, D.M., Petersen, S.E., Schlaggar, B.L., 2010. Prediction of individual brain maturity using fMRI. Science 329, 1358–1361. doi:10.1126/science.1194144

Feinberg, D.A., Moeller, S., Smith, S.M., Auerbach, E., Ramanna, S., Gunther, M., Glasser, M.F., Miller, K.L., Ugurbil, K., Yacoub, E., 2010. Multiplexed echo planar imaging for sub-second whole brain FMRI and fast diffusion imaging. PLoS ONE 5, e15710. doi:10.1371/journal.pone.0015710

Fernandes, B.S., Williams, L.M., Steiner, J., Leboyer, M., Carvalho, A.F., Berk, M., 2017. The new field of “precision psychiatry”. BMC Med. 15, 80. doi:10.1186/s12916-017-0849-x

Finn, E.S., Rosenberg, M.D., 2021. Beyond fingerprinting: Choosing predictive connectomes over reliable connectomes. Neuroimage 239, 118254. doi:10.1016/j.neuroimage.2021.118254

Finn, E.S., Scheinost, D., Finn, D.M., Shen, X., Papademetris, X., Constable, R.T., 2017. Can brain state be manipulated to emphasize individual differences in functional connectivity? Neuroimage 160, 140–151. doi:10.1016/j.neuroimage.2017.03.064

Finn, E.S., Shen, X., Scheinost, D., Rosenberg, M.D., Huang, J., Chun, M.M., Papademetris, X., Constable, R.T., 2015. Functional connectome fingerprinting: identifying individuals using patterns of brain connectivity. Nat. Neurosci. 18, 1664–1671. doi:10.1038/nn.4135

Fischl, B., 2012. FreeSurfer. Neuroimage 62, 774–781. doi:10.1016/j.neuroimage.2012.01.021

Gao, W., Elton, A., Zhu, H., Alcauter, S., Smith, J.K., Gilmore, J.H., Lin, W., 2014. Intersubject variability of and genetic effects on the brain’s functional connectivity during infancy. J. Neurosci. 34, 11288–11296. doi:10.1523/JNEUROSCI.5072-13.2014

Gaut, G., Turner, B., Lu, Z.-L., Li, X., Cunningham, W.A., Steyvers, M., 2019. Predicting Task and Subject Differences with Functional Connectivity and Blood-Oxygen-Level-Dependent Variability. Brain Connect. 9, 451–463. doi:10.1089/brain.2018.0632

Glasser, M.F., Sotiropoulos, S.N., Wilson, J.A., Coalson, T.S., Fischl, B., Andersson, J.L., Xu, J., Jbabdi, S., Webster, M., Polimeni, J.R., Van Essen, D.C., Jenkinson, M., WU-Minn HCP Consortium, 2013. The minimal preprocessing pipelines for the Human Connectome Project. Neuroimage 80, 105–124. doi:10.1016/j.neuroimage.2013.04.127

Golestani, A.M., Wei, L.L., Chen, J.J., 2016. Quantitative mapping of cerebrovascular reactivity using resting-state BOLD fMRI: Validation in healthy adults. Neuroimage 138, 147–163. doi:10.1016/j.neuroimage.2016.05.025

Greene, A.S., Gao, S., Scheinost, D., Constable, R.T., 2018. Task-induced brain state manipulation improves prediction of individual traits. Nat. Commun. 9, 2807. doi:10.1038/s41467-018-04920-3

Hansen, N., Ostermeier, A., 2001. Completely derandomized self-adaptation in evolution strategies. Evol. Comput. 9, 159–195. doi:10.1162/106365601750190398

Hastie, T., Tibshirani, R., Friedman, J., 2009. The Elements of Statistical Learning, second. ed.

Holiga, Š., Sambataro, F., Luzy, C., Greig, G., Sarkar, N., Renken, R.J., Marsman, J.-B.C., Schobel, S.A., Bertolino, A., Dukart, J., 2018. Test-retest reliability of task-based and resting-state blood oxygen level dependence and cerebral blood flow measures. PLoS ONE 13, e0206583. doi:10.1371/journal.pone.0206583

Jenkinson, M., Bannister, P., Brady, M., Smith, S., 2002. Improved optimization for the robust and accurate linear registration and motion correction of brain images. Neuroimage 17, 825–841. doi:10.1006/nimg.2002.1132

Jenkinson, M., Beckmann, C.F., Behrens, T.E., Woolrich, M.W., Smith, S.M., 2012. FSL. Neuroimage 62, 782–790. doi:10.1016/j.neuroimage.2011.09.015

Jiang, L., Zuo, X.-N., 2016. Regional homogeneity: A multimodal, multiscale neuroimaging marker of the human connectome. Neuroscientist 22, 486–505. doi:10.1177/1073858415595004

Jo, Y., Faskowitz, J., Esfahlani, F.Z., Sporns, O., Betzel, R.F., 2020. Subject identification using edge-centric functional connectivity. BioRxiv. doi:10.1101/2020.09.13.291898

Kaufmann, T., Alnæs, D., Brandt, C.L., Bettella, F., Djurovic, S., Andreassen, O.A., Westlye, L.T., 2018. Stability of the brain functional connectome fingerprint in individuals with schizophrenia. JAMA Psychiatry 75, 749–751. doi:10.1001/jamapsychiatry.2018.0844

Kaufmann, T., Alnæs, D., Doan, N.T., Brandt, C.L., Andreassen, O.A., Westlye, L.T., 2017. Delayed stabilization and individualization in connectome development are related to psychiatric disorders. Nat. Neurosci. 20, 513–515. doi:10.1038/nn.4511

Kong, R., Li, J., Orban, C., Sabuncu, M.R., Liu, H., Schaefer, A., Sun, N., Zuo, X.-N., Holmes, A.J., Eickhoff, S.B., Yeo, B.T.T., 2019. Spatial Topography of Individual-Specific Cortical Networks Predicts Human Cognition, Personality, and Emotion. Cereb. Cortex 29, 2533–2551. doi:10.1093/cercor/bhy123

Kumar, K., Toews, M., Chauvin, L., Colliot, O., Desrosiers, C., 2018. Multi-modal brain fingerprinting: A manifold approximation based framework. Neuroimage 183, 212–226. doi:10.1016/j.neuroimage.2018.08.006

Laumann, T.O., Gordon, E.M., Adeyemo, B., Snyder, A.Z., Joo, S.J., Chen, M.-Y., Gilmore, A.W., McDermott, K.B., Nelson, S.M., Dosenbach, N.U.F., Schlaggar, B.L., Mumford, J.A., Poldrack, R.A., Petersen, S.E., 2015. Functional system and areal organization of a highly sampled individual human brain. Neuron 87, 657–670. doi:10.1016/j.neuron.2015.06.037

Li, C., Yang, G., Li, M., Li, B., 2018. Fluid intelligence relates to the resting state amplitude of low-frequency fluctuation and functional connectivity: a multivariate pattern analysis. Neuroreport 29, 8–12. doi:10.1097/WNR.0000000000000917

Li, K., Wisner, K., Atluri, G., 2021. Feature selection framework for functional connectome fingerprinting. Human Brain Mapping.

Liégeois, R., Laumann, T.O., Snyder, A.Z., Zhou, J., Yeo, B.T.T., 2017. Interpreting temporal fluctuations in resting-state functional connectivity MRI. Neuroimage 163, 437–455. doi:10.1016/j.neuroimage.2017.09.012

Liu, J., Liao, X., Xia, M., He, Y., 2018. Chronnectome fingerprinting: Identifying individuals and predicting higher cognitive functions using dynamic brain connectivity patterns. Hum. Brain Mapp. 39, 902–915. doi:10.1002/hbm.23890

Mansour L, S., Tian, Y., Yeo, B.T.T., Cropley, V., Zalesky, A., 2021. High-resolution connectomic fingerprints: Mapping neural identity and behavior. Neuroimage 229, 117695. doi:10.1016/j.neuroimage.2020.117695

Mantwill, M., Gell, M., Krohn, S., Finke, C., 2021. Fingerprinting and behavioural prediction rest on distinct functional systems of the human connectome. BioRxiv. doi:10.1101/2021.02.07.429922

Miranda-Dominguez, O., Mills, B.D., Carpenter, S.D., Grant, K.A., Kroenke, C.D., Nigg, J.T., Fair, D.A., 2014. Connectotyping: model based fingerprinting of the functional connectome. PLoS ONE 9, e111048. doi:10.1371/journal.pone.0111048

Moeller, S., Yacoub, E., Olman, C.A., Auerbach, E., Strupp, J., Harel, N., Uğurbil, K., 2010. Multiband multislice GE-EPI at 7 tesla, with 16-fold acceleration using partial parallel imaging with application to high spatial and temporal whole-brain fMRI. Magn. Reson. Med. 63, 1144–1153. doi:10.1002/mrm.22361

More, S., Eickhoff, S.B., Caspers, J., Patil, K.R., 2021. Confound removal and normalization in practice: A neuroimaging based sex prediction case study, in: Dong, Y., Ifrim, G., Mladenić, D., Saunders, C., Van Hoecke, S. (Eds.), ECML PKDD 2020: Demo Track, Lecture Notes in Computer Science. Springer International Publishing, Ghent, Belgium, pp. 3–18. doi:10.1007/978-3-030-67670-4_1

Mueller, S., Wang, D., Fox, M.D., Yeo, B.T.T., Sepulcre, J., Sabuncu, M.R., Shafee, R., Lu, J., Liu, H., 2013. Individual variability in functional connectivity architecture of the human brain. Neuron 77, 586–595. doi:10.1016/j.neuron.2012.12.028

Ni, H., Song, M., Qin, J., Jiang, T., 2021. Individual discriminative ability of resting functional brain connectivity is susceptible to the time span of MRI scans. Neuroscience. doi:10.1016/j.neuroscience.2021.12.008

Noble, S., Scheinost, D., Constable, R.T., 2019. A decade of test-retest reliability of functional connectivity: A systematic review and meta-analysis. Neuroimage 203, 116157. doi:10.1016/j.neuroimage.2019.116157

Noble, S., Spann, M.N., Tokoglu, F., Shen, X., Constable, R.T., Scheinost, D., 2017. Influences on the Test-Retest Reliability of Functional Connectivity MRI and its Relationship with Behavioral Utility. Cereb. Cortex 27, 5415–5429. doi:10.1093/cercor/bhx230

Nooner, K.B., Colcombe, S.J., Tobe, R.H., Mennes, M., Benedict, M.M., Moreno, A.L., Panek, L.J., Brown, S., Zavitz, S.T., Li, Q., Sikka, S., Gutman, D., Bangaru, S., Schlachter, R.T., Kamiel, S.M., Anwar, A.R., Hinz, C.M., Kaplan, M.S., Rachlin, A.B., Adelsberg, S., Milham, M.P., 2012. The NKI-Rockland Sample: A Model for Accelerating the Pace of Discovery Science in Psychiatry. Front. Neurosci. 6, 152. doi:10.3389/fnins.2012.00152

Nostro, A.D., Müller, V.I., Varikuti, D.P., Pläschke, R.N., Hoffstaedter, F., Langner, R., Patil, K. R., Eickhoff, S.B., 2018. Predicting personality from network-based resting-state functional connectivity. Brain Struct. Funct. 223, 2699–2719. doi:10.1007/s00429-018-1651-z

Pinto, J., Bright, M.G., Bulte, D.P., Figueiredo, P., 2020. Cerebrovascular reactivity mapping without gas challenges: A methodological guide. Front. Physiol. 11, 608475. doi:10.3389/fphys.2020.608475

Power, J.D., Cohen, A.L., Nelson, S.M., Wig, G.S., Barnes, K.A., Church, J.A., Vogel, A.C., Laumann, T.O., Miezin, F.M., Schlaggar, B.L., Petersen, S.E., 2011. Functional network organization of the human brain. Neuron 72, 665–678. doi:10.1016/j.neuron.2011.09.006

Reinen, J.M., Chén, O.Y., Hutchison, R.M., Yeo, B.T.T., Anderson, K.M., Sabuncu, M.R., Öngür, D., Roffman, J.L., Smoller, J.W., Baker, J.T., Holmes, A.J., 2018. The human cortex possesses a reconfigurable dynamic network architecture that is disrupted in psychosis. Nat. Commun. 9, 1157. doi:10.1038/s41467-018-03462-y

Repovš, G., Barch, D.M., 2012. Working memory related brain network connectivity in individuals with schizophrenia and their siblings. Front. Hum. Neurosci. 6, 137. doi:10.3389/fnhum.2012.00137

Schaefer, A., Kong, R., Gordon, E.M., Laumann, T.O., Zuo, X.-N., Holmes, A.J., Eickhoff, S.B., Yeo, B.T.T., 2018. Local-Global Parcellation of the Human Cerebral Cortex from Intrinsic Functional Connectivity MRI. Cereb. Cortex 28, 3095–3114. doi:10.1093/cercor/bhx179

Schultz, D.H., Cole, M.W., 2016. Higher Intelligence Is Associated with Less Task-Related Brain Network Reconfiguration. J. Neurosci. 36, 8551–8561. doi:10.1523/JNEUROSCI.0358-16.2016

Seitzman, B.A., Gratton, C., Laumann, T.O., Gordon, E.M., Adeyemo, B., Dworetsky, A., Kraus, B.T., Gilmore, A.W., Berg, J.J., Ortega, M., Nguyen, A., Greene, D.J., McDermott, K.B., Nelson, S.M., Lessov-Schlaggar, C.N., Schlaggar, B.L., Dosenbach, N.U.F., Petersen, S.E., 2019. Trait-like variants in human functional brain networks. Proc Natl Acad Sci USA 116, 22851–22861. doi:10.1073/pnas.1902932116

Setsompop, K., Gagoski, B.A., Polimeni, J.R., Witzel, T., Wedeen, V.J., Wald, L.L., 2012. Blipped-controlled aliasing in parallel imaging for simultaneous multislice echo planar imaging with reduced g-factor penalty. Magn. Reson. Med. 67, 1210–1224. doi:10.1002/mrm.23097

Shen, X., Tokoglu, F., Papademetris, X., Constable, R.T., 2013. Groupwise whole-brain parcellation from resting-state fMRI data for network node identification. Neuroimage 82, 403–415. doi:10.1016/j.neuroimage.2013.05.081

Shukla, D.K., Keehn, B., Müller, R.A., 2010. Regional homogeneity of fMRI time series in autism spectrum disorders. Neurosci. Lett. 476, 46–51. doi:10.1016/j.neulet.2010.03.080

Thiele, J.A., Faskowitz, J., Sporns, O., Hilger, K., 2021. Multi-Task Brain Network Reconfiguration is Inversely Associated with Human Intelligence. BioRxiv. doi:10.1101/2021.07.31.454563

Tipnis, U., Abbas, K., Tran, E., Amico, E., Li, S., Alan, 2020. Data Release and Assessment on Brain Fingerprints.

Tomasi, D., Wang, G.-J., Volkow, N.D., 2013. Energetic cost of brain functional connectivity. Proc Natl Acad Sci USA 110, 13642–13647. doi:10.1073/pnas.1303346110

Van Essen, D.C., Smith, S.M., Barch, D.M., Behrens, T.E.J., Yacoub, E., Ugurbil, K., WU-Minn HCP Consortium, 2013. The WU-Minn Human Connectome Project: an overview. Neuroimage 80, 62–79. doi:10.1016/j.neuroimage.2013.05.041

Wang, L., Song, M., Jiang, T., Zhang, Y., Yu, C., 2011. Regional homogeneity of the restingstate brain activity correlates with individual intelligence. Neurosci. Lett. 488, 275–278. doi:10.1016/j.neulet.2010.11.046

Wang, P., Yang, J., Yin, Z., Duan, J., Zhang, R., Sun, J., Xu, Y., Liu, L., Chen, X., Li, H., Kang, J., Zhu, Y., Deng, X., Chang, M., Wei, S., Zhou, Y., Jiang, X., Wang, F., Tang, Y., 2019. Amplitude of low-frequency fluctuation (ALFF) may be associated with cognitive impairment in schizophrenia: a correlation study. BMC Psychiatry 19, 30. doi:10.1186/s12888-018-1992-4

Woolrich, M.W., Ripley, B.D., Brady, M., Smith, S.M., 2001. Temporal autocorrelation in univariate linear modeling of FMRI data. Neuroimage 14, 1370–1386. doi:10.1006/nimg.2001.0931

Xifra-Porxas, A., Kassinopoulos, M., Mitsis, G.D., 2021. Physiological and motion signatures in static and time-varying functional connectivity and their subject identifiability. eLife 10. doi:10.7554/eLife.62324

Xu, J., Moeller, S., Strupp, J., Auerbach, E., Feinberg, D.A., Ugurbil, K., Yacoub, E., 2021. Highly Accelerated Whole Brain Imaging Using Aligned-Blipped-Controlled-Aliasing Multiband EPI. Proc. Int. Soc. Mag. Reson. Med. 20, 2306.

Xu, T., Opitz, A., Craddock, R.C., Wright, M.J., Zuo, X.-N., Milham, M.P., 2016. Assessing variations in areal organization for the intrinsic brain: from fingerprints to reliability. Cereb. Cortex 26, 4192–4211. doi:10.1093/cercor/bhw241

Yeo, B.T.T., Krienen, F.M., Sepulcre, J., Sabuncu, M.R., Lashkari, D., Hollinshead, M., Roffman, J.L., Smoller, J.W., Zöllei, L., Polimeni, J.R., Fischl, B., Liu, H., Buckner, R.L., 2011. The organization of the human cerebral cortex estimated by intrinsic functional connectivity. J. Neurophysiol. 106, 1125–1165. doi:10.1152/jn.00338.2011

Yuan, R., Di, X., Kim, E.H., Barik, S., Rypma, B., Biswal, B.B., 2013. Regional homogeneity of resting-state fMRI contributes to both neurovascular and task activation variations. Magn. Reson. Imaging 31, 1492–1500. doi:10.1016/j.mri.2013.07.005

Zang, Y., Jiang, T., Lu, Y., He, Y., Tian, L., 2004. Regional homogeneity approach to fMRI data analysis. Neuroimage 22, 394–400. doi:10.1016/j.neuroimage.2003.12.030

Zang, Y.-F., He, Y., Zhu, C.-Z., Cao, Q.-J., Sui, M.-Q., Liang, M., Tian, L.-X., Jiang, T.-Z., Wang, Y.-F., 2007. Altered baseline brain activity in children with ADHD revealed by resting-state functional MRI. Brain Dev. 29, 83–91. doi:10.1016/j.braindev.2006.07.002

Zhang, S., Spoletini, L.J., Gold, B.P., Morgan, V.L., Rogers, B.P., Chang, C., 2021. Interindividual Signatures of fMRI Temporal Fluctuations. Cereb. Cortex 31, 4450–4463. doi:10.1093/cercor/bhab099

Zhao, X., Yao, J., Lv, Y., Zhang, X., Han, C., Chen, L., Ren, F., Jin, Z., Li, Y., Sui, Y., 2019. Abnormalities of regional homogeneity and its correlation with clinical symptoms in Naïve patients with first-episode schizophrenia. Brain Imaging Behav. 13, 503–513. doi:10.1007/s11682-018-9882-4

Zou, Q.-H., Zhu, C.-Z., Yang, Y., Zuo, X.-N., Long, X.-Y., Cao, Q.-J., Wang, Y.-F., Zang, Y.-F., 2008. An improved approach to detection of amplitude of low-frequency fluctuation (ALFF) for resting-state fMRI: fractional ALFF. J. Neurosci. Methods 172, 137–141. doi:10.1016/j.jneumeth.2008.04.012

Zuo, X.-N., Di Martino, A., Kelly, C., Shehzad, Z.E., Gee, D.G., Klein, D.F., Castellanos, F.X., Biswal, B.B., Milham, M.P., 2010. The oscillating brain: complex and reliable. Neuroimage 49, 1432–1445. doi:10.1016/j.neuroimage.2009.09.037

