## Supplementary Materials for "Spatially localized fMRI metrics as predictive and highly distinct state-independent fingerprints"

#### Data

##### OpenNeuro.ds000115 Working memory in healthy and schizophrenic individuals

We examined the OpenNeuro.ds000115 "Working memory in healthy and schizophrenic individuals" dataset (Repovš and Barch, 2012). Subjects were included when data was available of the 0- and 2-back fMRI tasks and intelligence scores. This resulted in the inclusion of 19 patients with schizophrenia (mean age = 23.9, SD = 3.7, 6 females) and 20 healthy controls (HC; mean age = 20.6, SD = 5.1, 8 females). The fMRI data of the 0-back and 2-back tasks were used for analyses. The WAIS Matrix Reasoning (WAIS_MATRICS_SCALE) and Vocabulary subtest scores (WAIS_VOCAB_SCALED) were used for investigating the correlations between fingerprint stability and intelligence.

##### Human Connectome Project Early Psychosis sample

We examined data of the HCP Early Psychosis Release 1.1 (https://www.humanconnectome.org/study/human-connectome-project-for-early-psychosis) which includes minimally preprocessed structural data and unpreprocessed fMRI data of 161 subjects. This dataset includes individuals within the first five years of the onset of psychotic symptoms. Given that the subgroup of patients with affective psychosis only included 23 subjects, we focused on the individuals with non-affective psychosis and healthy controls. Furthermore, we selected subjects with data available of 1) all four resting state scans, 2) fluid and crystallized cognition composite scores, and 3) age and sex. This resulted in the inclusion of 53 healthy controls and 70 individuals with non-affective psychosis, diagnosed with schizophrenia, schizophreniform disorder, schizoaffective disorder, psychosis NOS, delusional disorder, or brief psychotic disorder.

##### Enhanced Nathan Kline Institute (eNKI) sample

Data of 450 clinically healthy subjects (i.e. without a psychiatric or neurological diagnosis or use of medication for the treatment of any such conditions) from the community-based sample of the enhanced Nathan Kline Institute dataset were included (mean age = 52.1, SD = 16.7, 306 females) (Nooner et al., 2012). This subset was selected based on the availability of both resting-state and breath-holding scans, with the exclusion of subjects with excessive movement or who fell asleep during the resting-state scanning.

### Methods

#### Figures

All figures of identification accuracies were created with R version 3.6.1 (R Core Team 2019) using packages reshape2 (Wickham, 2007), ggplot2 (Wickham, 2016), extrafont (Chang, 2014), and grid.

#### Effect of sphere size around Power coordinates on identification accuracies - smaller regions of interest (ROIs) yield higher identification accuracies

The effect of sphere size on identification accuracies was investigated, as sphere size is known to affect the calculation of regional homogeneity (ReHo) (Zang et al., 2004). We tested spheres with radiuses of 3, 5, and 10 mm.

#### Replication and additional analyses sample Finn et al. (2015)

To replicate the results shown in Finn et al. (2015), the same sample was included (Finn et al., 2015). One subject was not included as their data was not available (anymore), leaving 125 subjects for analyses. Of these 125 subjects, 47 subjects overlapped with our large HCP sample (n=399). Data from the minimally preprocessed Human Connectome Project (HCP) pipeline of the same sessions reported by Finn et al. (2015) was included. For the resting-state fMRI data, this did not include the removal of noise components as obtained with ICA FIX. Additional preprocessing and calculation of connectivity matrices were done in Matlab 2019b (Mathworks Inc, Natick, MA). This included the removal of 12 motion parameters (provided by HCP as Movement_Regressors_dt.txt files), regression of the mean time courses of WM, CSF, and global signal, removal of the polynomial trend up to order 3, and low pass filtering (FFT; 0-0.26 Hz). No smoothing was applied. The brain was parcellated into 268 regions according to the Shen parcellation (Shen et al., 2013). Pearson correlation coefficients were calculated between the mean time courses of each pair of regions, resulting in 268 × 268 symmetric FC matrices for each session for each subject. This was done separately for the LR- and RL-phase encoding runs, and the matrices were consequently averaged. Each element in the resulting matrices (i.e. 125 subjects x 6 sessions) reflects the connectivity between two regions. Next, identification accuracy was calculated. In addition to the FC analyses described in Finn et al. (2015), we also calculated identification accuracies for ReHo, ALFF, and fALFF, using the same sample and preprocessing parameters. For comparison, we also calculated identification accuracies for FC, ReHo, ALFF, and fALFF with three other parcellations (i.e. Power (3-mm spheres), Dosenbach and Schaefer).

#### Local fingerprints in two clinical samples

We examined the OpenNeuro.ds000115 Working memory in healthy and schizophrenic individuals' dataset. The sample is described in the Data section. The data were preprocessed with fMRIPrep version 1.5.9, which has been described in detail elsewhere (Esteban et al., 2019). Briefly, for the structural MRI images, this included intensity non-uniformity (INU) correction, skull-stripping of the T1w image, followed by tissue segmentation, and spatial normalization to MNI-space using nonlinear registration. For the resting state and breath-holding data, a reference image and its skull-stripped version were first generated. Next, head motion relative to the reference image was estimated and slice-time correction was performed. The BOLD images were then coregistered to the native space of the subjects' T1w image, resampled to correct for motion, and mapped to MNI space. Note that ICA-AROMA denoising was not used as this smooths the data resulting in more spatial homogeneity and consequently more uniform and non-specific ReHo estimates. Last, mean cerebrospinal fluid and white matter time courses, as well as 12 motion parameters, were regressed out and the data were bandpass filtered (0.01-0.1Hz). Regions were created with 3-mm spheres around the Power coordinates. Identification accuracy was calculated as described in the Methods section of the main text.

Additionally, we examined the HCP Early Psychosis dataset. The sample is described in the Data section. We used the HCP pipelines to minimally preprocess this data (Glasser et al., 2013) (script GenericfMRIVolumeProcessingPipeline) and apply multi-run ICA-FIX cleaning (script IcaFixProcessing). The volumetric time series of four resting state scans (resting state 1 and 2 both acquired with AP and PA phase encoding) were used for analyses. Mean CSF and WM time courses, as well as 12 motion parameters, were regressed out and the data were bandpass filtered (0.01-0.1Hz). Regions were created with 3-mm spheres around the Power coordinates. Next, the fingerprints were calculated for both AP and PA phase-encoding runs and consequently averaged. Identification accuracy was calculated as described in the Methods section of the main text. As the number of patients was higher than the number of healthy controls, we randomly selected a subsample of 53 patients to calculate identification accuracies and the correlations between fingerprint stability and intelligence. This was repeated 100 times and the mean across repetitions was used. The fluid and crystallized intelligence composite scores (nih_fluidcogcomp_unadjusted and nih_crycogcomp_unadjusted; age-adjusted intelligence scores were missing for many subjects) were used for investigations of the partial correlation with intelligence controlling for age.

#### Signals underlying regional homogeneity

The selection of data of the Enhanced Nathan Kline Institute (eNKI) sample is described in the Data section. Next, the data were preprocessed with fMRIPrep version 20.1.1 which has been described in detail elsewhere (Esteban et al., 2019). Briefly, for the structural MRI images, this included intensity non-uniformity (INU) correction, skull-stripping of the T1w image, followed by tissue segmentation, and spatial normalization to MNI-space using nonlinear registration. For the resting state and breath-holding data, a reference image and its skull-stripped version were first generated. Next, head motion relative to the reference image was estimated and slice-time correction was performed. The BOLD images were then coregistered to the native space of the subjects' T1w image, resampled to correct for motion, and mapped to MNI space. Note that ICA-AROMA denoising was not used as this smooths the data resulting in more spatial homogeneity and consequently more uniform and non-specific ReHo estimates. Lastly, mean cerebrospinal fluid and white matter time courses, as well as 12 motion parameters, were regressed out and the data were bandpass filtered (0.01-0.1Hz). For the breath-holding task, fingerprints were calculated only including the breath-holding (BH) parts of the task (i.e. start 18 seconds after task onset, lasting for 18 seconds, alternating with 18 seconds of breathing in and out). Regions were created with 3-mm spheres around the Power coordinates. Identification accuracy was calculated as described in the Methods section of the main text.

### Results

#### Effect of sphere size around Power coordinates on identification accuracies - smaller ROIs yield higher identification accuracy

We found that sphere sizes of 3 mm yielded the highest identification accuracy for all the fingerprints (Fig. S1). The main results are therefore reported with spheres of size 3 mm.

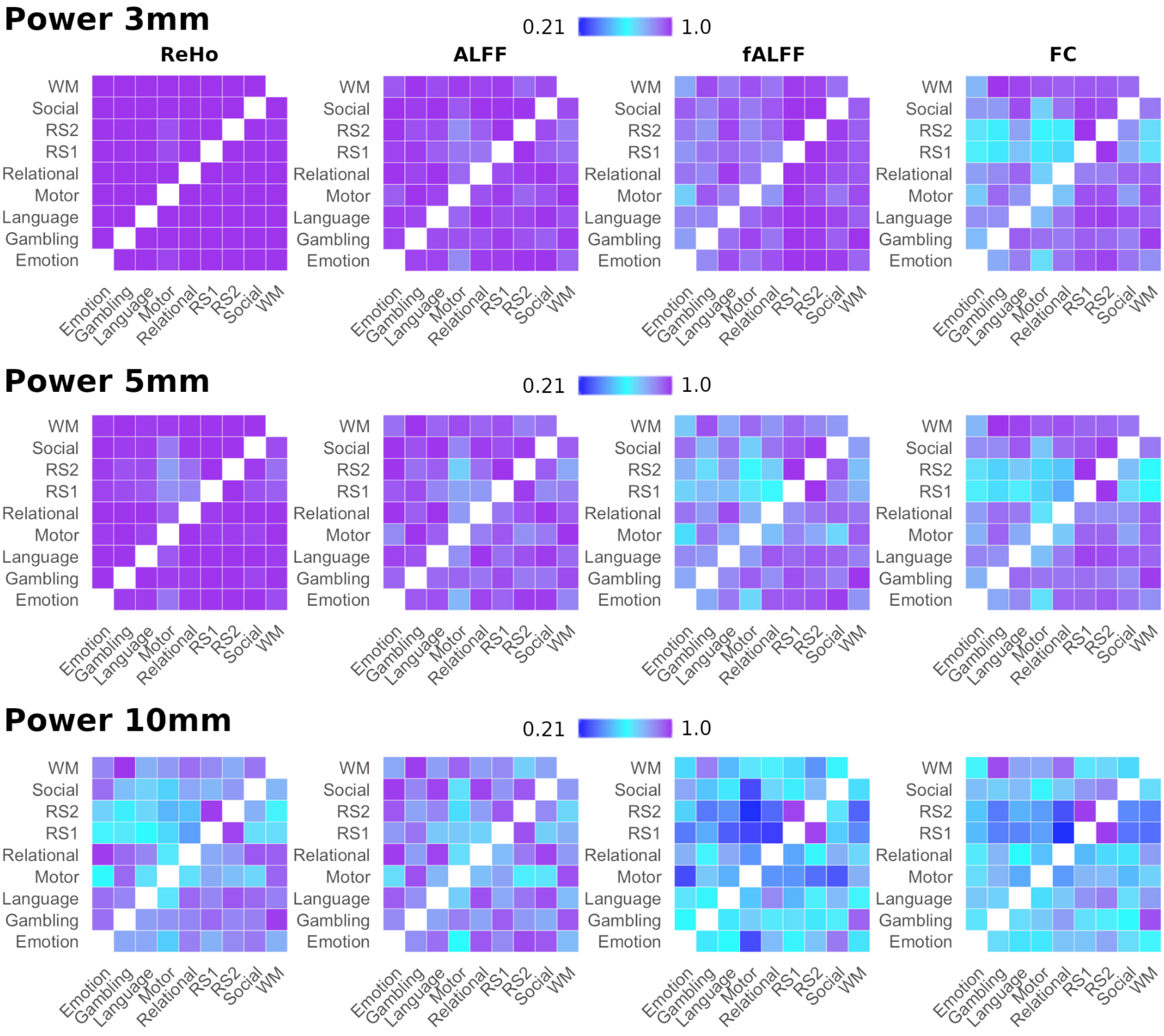

**Fig. S1**: comparison of identification accuracies for several functional fingerprints with different sphere sizes around the Power coordinates in the Human Connectome Project sample. The highest identification accuracies were seen for 3-mm sphere sizes, followed by 5-mm and 10-mm. Therefore, the 3-mm spheres were used for the main analyses. Abbreviations: ReHo = regional homogeneity; ALFF = Amplitude of Low Frequency Fluctuations; fALFF = fractional Amplitude of Low Frequency Fluctuations; FC = functional connectome; RS = resting state.

### Idiff

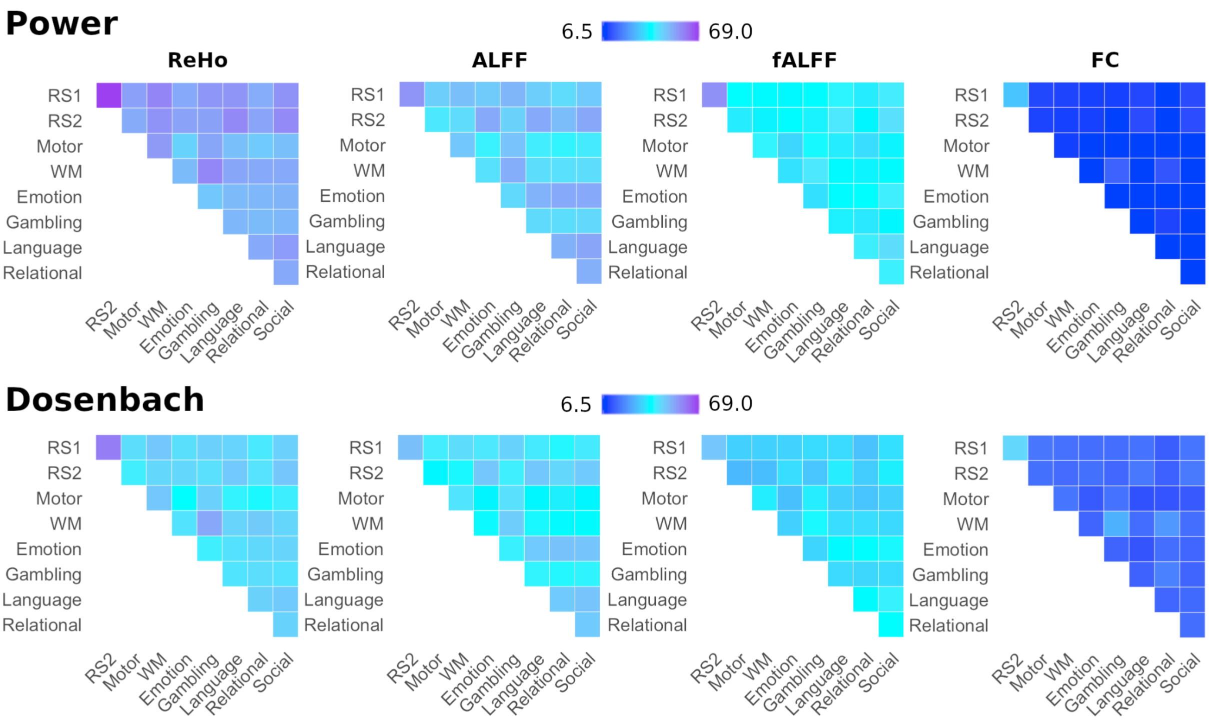


**Fig. S2**: comparison of Idiff of functional fingerprints in the Human Connectome Project sample. Abbreviations: ReHo = regional homogeneity; ALFF = Amplitude of Low Frequency Fluctuations; fALFF = fractional Amplitude of Low Frequency Fluctuations; FC = functional connectome; RS = resting state.

### Fingerprint identification accuracies with Dosenbach, Shen, and Schaefer parcellations

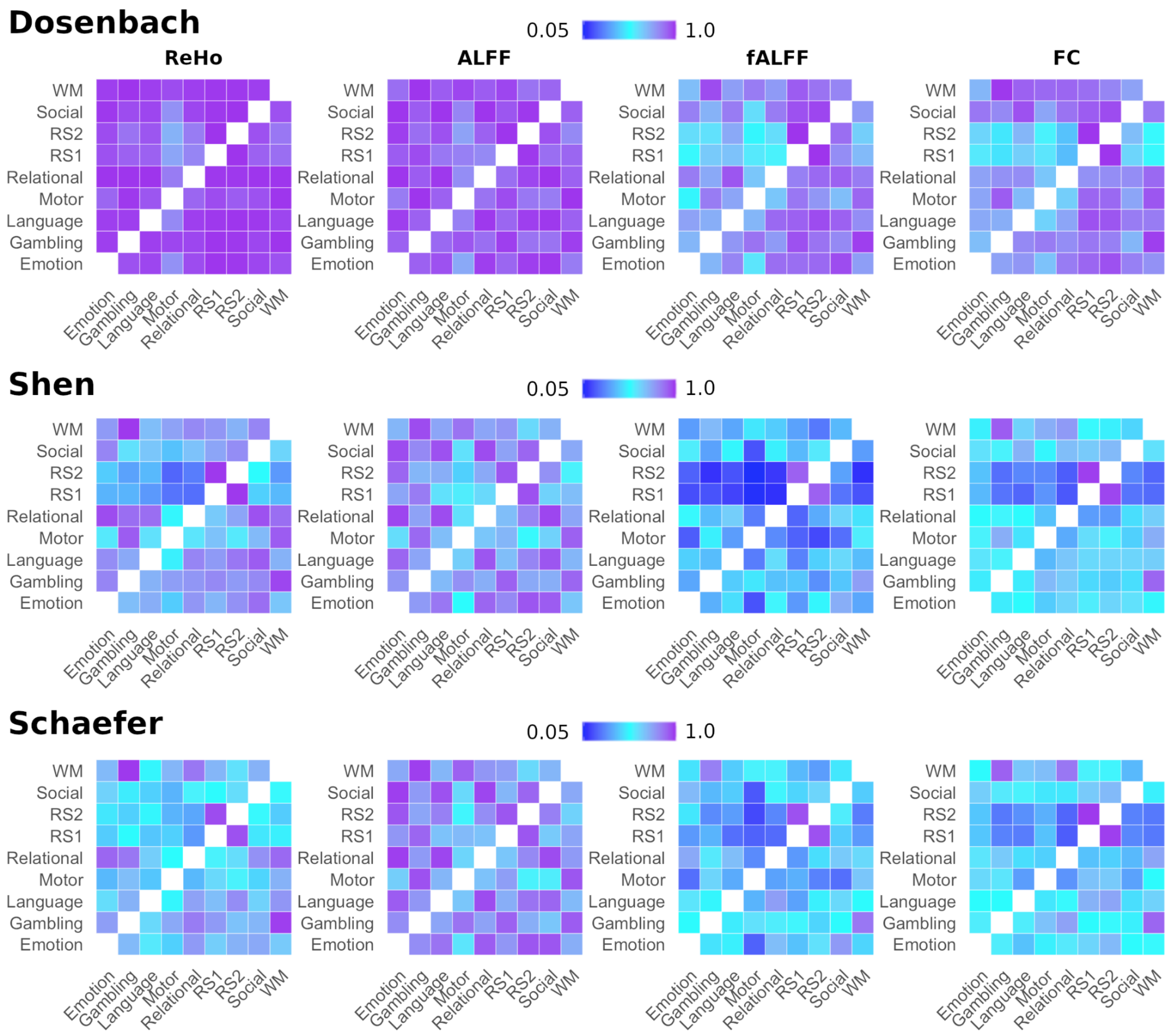


**Fig. S3**: comparison of identification accuracies of local and global functional fingerprints in the Human Connectome Project sample for the Dosenbach, Shen, and Schaefer 300 parcellations. Abbreviations: ReHo = regional homogeneity; ALFF = Amplitude of Low Frequency Fluctuations; fALFF = fractional Amplitude of Low Frequency Fluctuations; FC = functional connectome; RS = resting state.

#### Identification accuracies of functional fingerprints in the HCP sample after confound removal

After confound removal, identification accuracies were still highest for ReHo (Fig. S4). In contrast with the results without confound regression, this was followed by FC, fALFF, and ALFF (in contrast to ReHo-ALFF-fALFF-FC). These results suggest a larger influence of confounders on global fingerprints such as the FC.

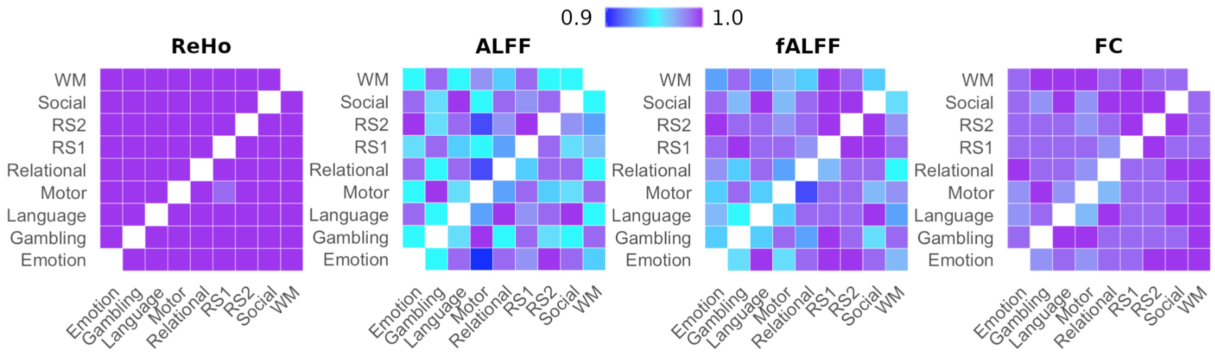


**Fig. S4:** comparison of identification accuracies of functional fingerprints in the Human Connectome Project sample after confound removal (i.e. sex, age, and intracranial volume). Abbreviations: ReHo = regional homogeneity; ALFF = Amplitude of Low Frequency Fluctuations; fALFF = fractional Amplitude of Low Frequency Fluctuations; FC = functional connectome; RS = resting state.

### Replication and additional analyses sample Finn et al. (2015)

First, we replicated the results reported by Finn et al. (2015) (Fig. S5). Finn et al. (2015) reported a mean identification accuracy of 94.4% between the FCs of two resting-state fMRI sessions. We show a mean accuracy of 90% (Fig. S5), which is in line with another replication (91%) (Venkatesh et al., 2020). Accuracies between FCs of task-based fMRI sessions and between resting-state and task-based fMRI sessions were also similar (i.e. 51-80%) to results reported in Finn et al. (2015) (i.e. > 54%).

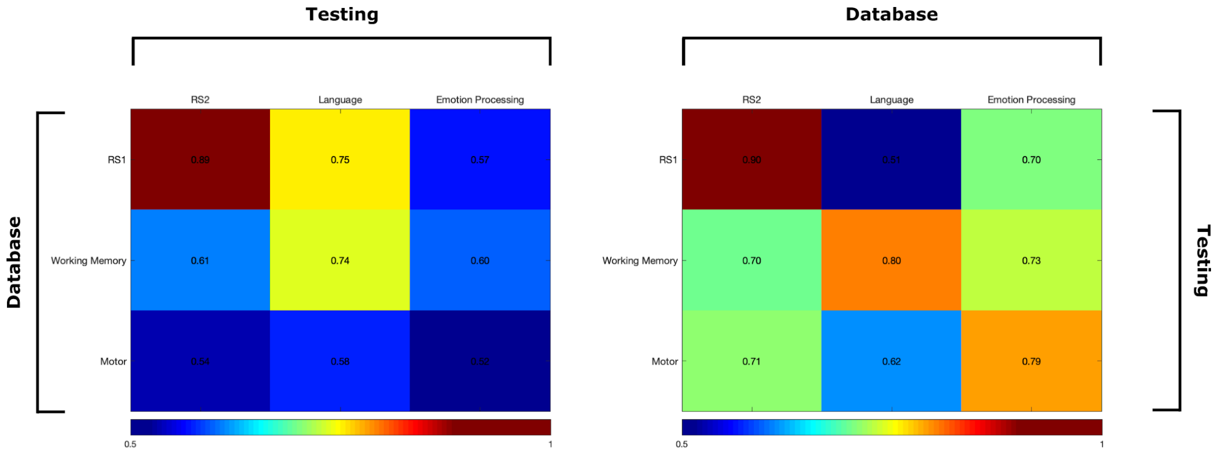


**Fig. S5**: comparison of identification accuracies of functional connectome (FC) fingerprints based on Pearson correlations between time courses for 125 subjects of the Human Connectome Project sample (replication of (Finn et al., 2015)). Abbreviations: RS = resting state.

Second, we calculated identification accuracies for the FC, as well as the local fingerprints ReHo, ALFF, and fALFF using four different parcellations (Fig. S6). For the Power spheres (3 mm) and Dosenbach parcellation, similarly to the results in the large HCP sample, we found near-perfect identification accuracies for ReHo, followed by ALFF, the FC, and fALFF. One difference was observed compared to results in the large HCP sample:

identification accuracy was higher for the FC than for fALFF. Results for the Schaefer parcellation were the same as in the large HCP sample: highest identification accuracies for ALFF, followed by ReHo, FC, and fALFF. Results for the Shen parcellation were slightly different compared to the large HCP-sample: highest identification accuracies for ReHo, followed by ALFF, FC, and fALFF (versus ALFF, ReHo, FC, fALFF in the large HCP-sample).

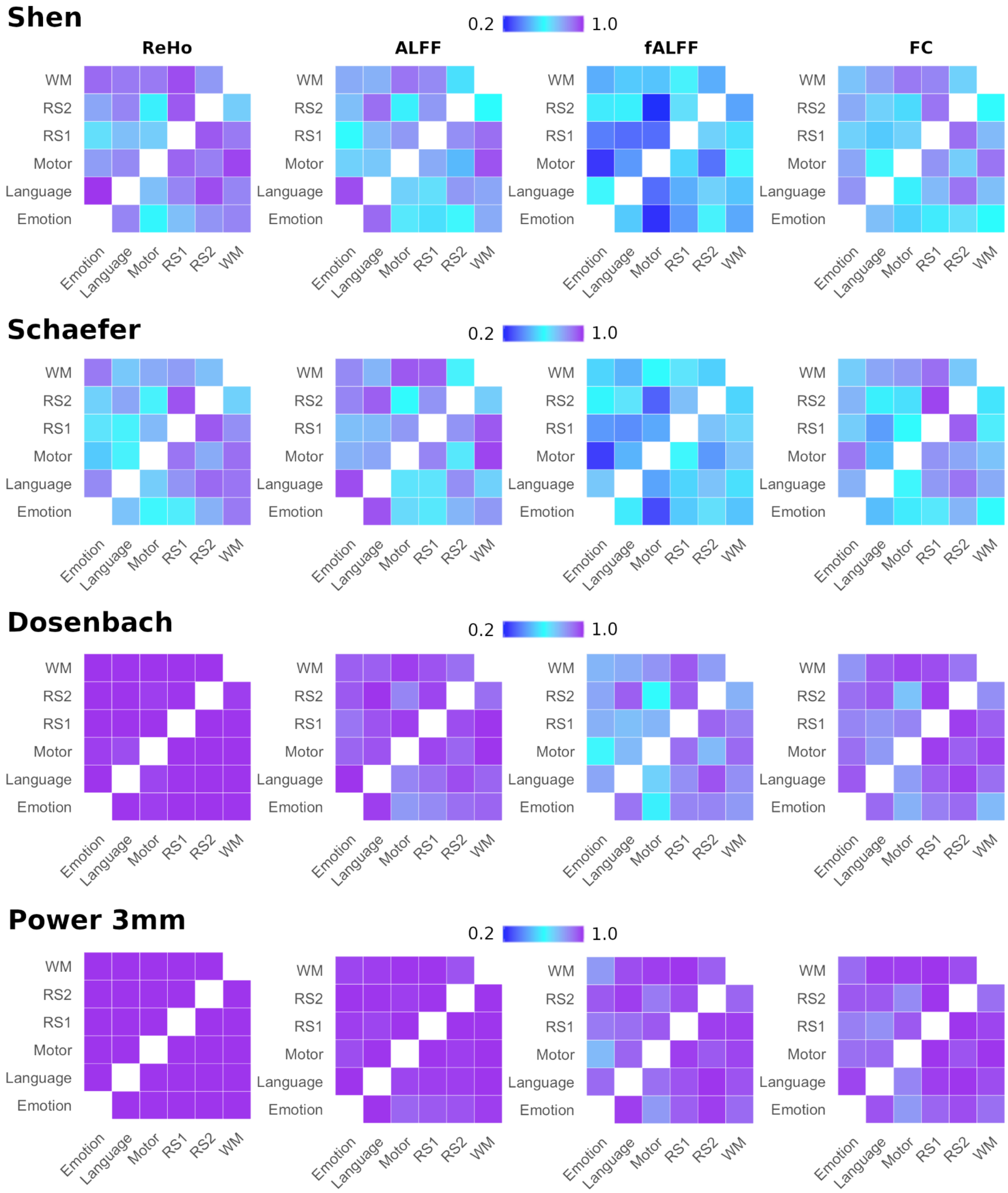


**Fig. S6**: comparison of identification accuracies of the FC with other functional fingerprints for 125 subjects of the Human Connectome Project sample. The same sample was included and similar preprocessing parameters were used as reported in Finn et al. (2015). Abbreviations: ReHo = regional homogeneity; ALFF = Amplitude of Low Frequency

Fluctuations; fALFF = fractional Amplitude of Low Frequency Fluctuations; FC = functional connectome; RS = resting state.

### Fingerprints predict intelligence

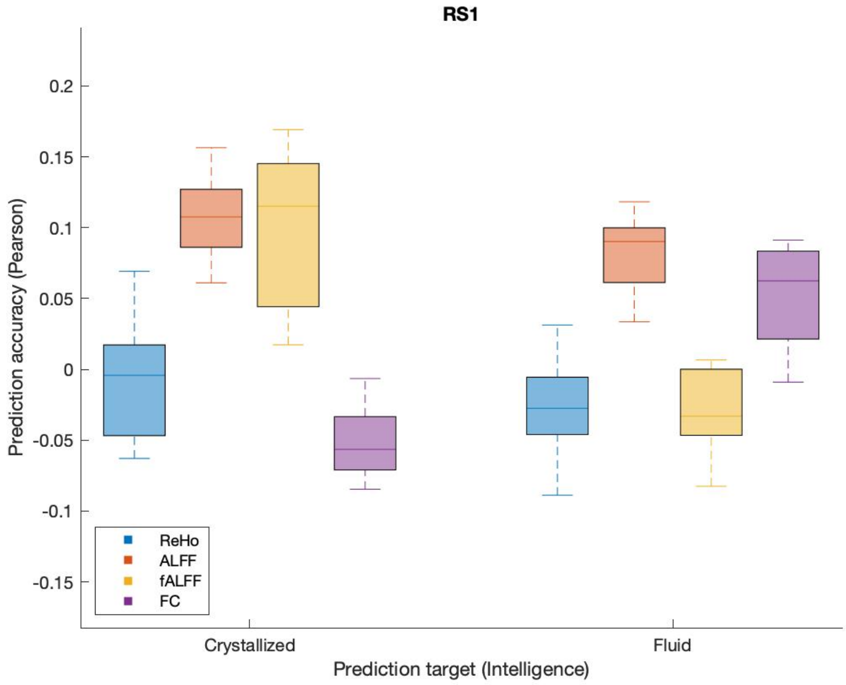


**Fig. S7**: Prediction of crystallized and fluid intelligence using the fingerprints of the Resting State 1 fMRI session of the Human Connectome Project sample. Each boxplot shows the prediction performance across five cross-validation runs.

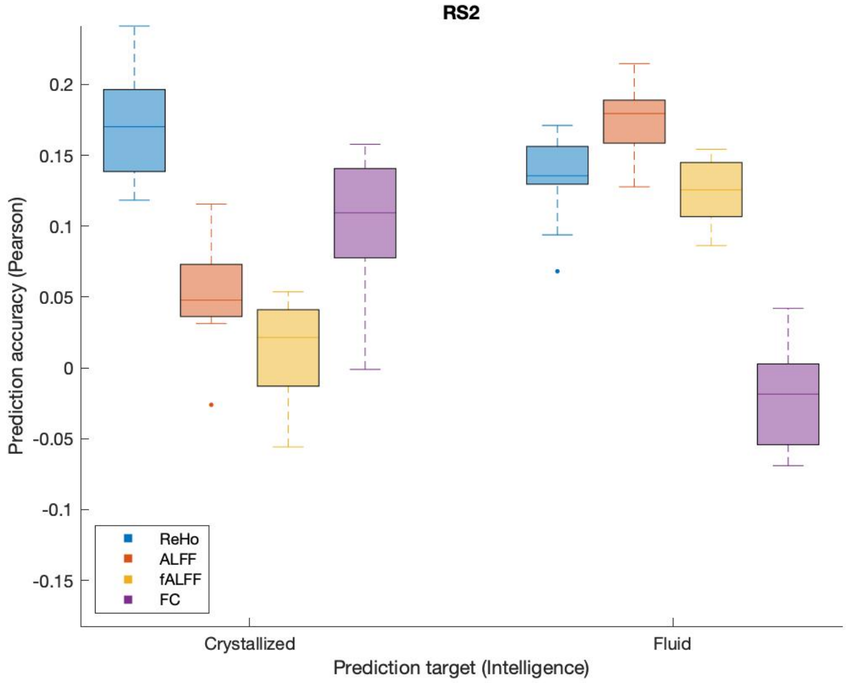

**Fig. S8**: Prediction of crystallized and fluid intelligence using the fingerprints of the Resting State 2 fMRI session of the Human Connectome Project sample. Each boxplot shows the prediction performance across five cross-validation runs.

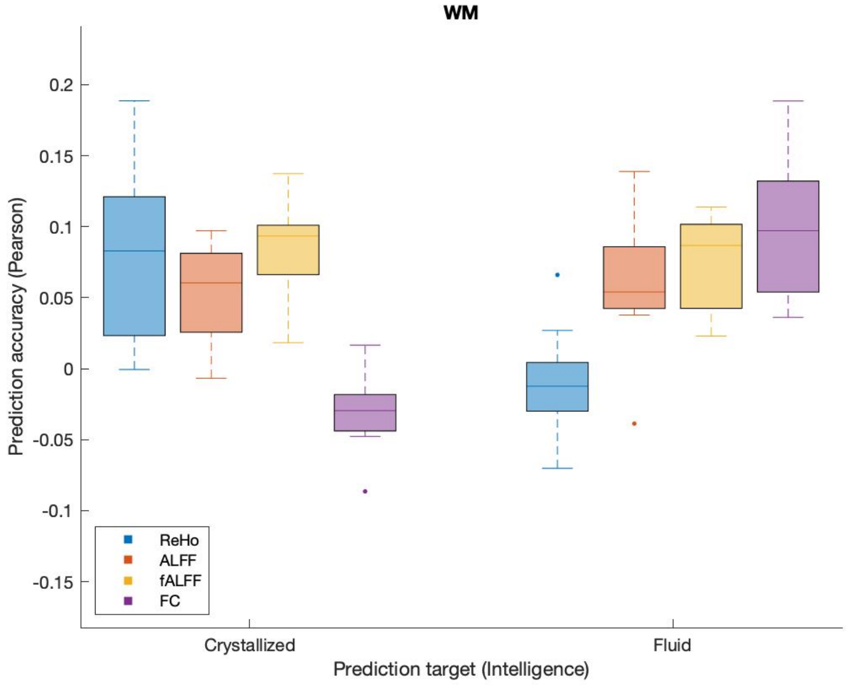


**Fig. S9:** Prediction of crystallized and fluid intelligence using the fingerprints of the working memory (WM) fMRI task of the Human Connectome Project sample. Each boxplot shows the prediction performance across five cross-validation runs.

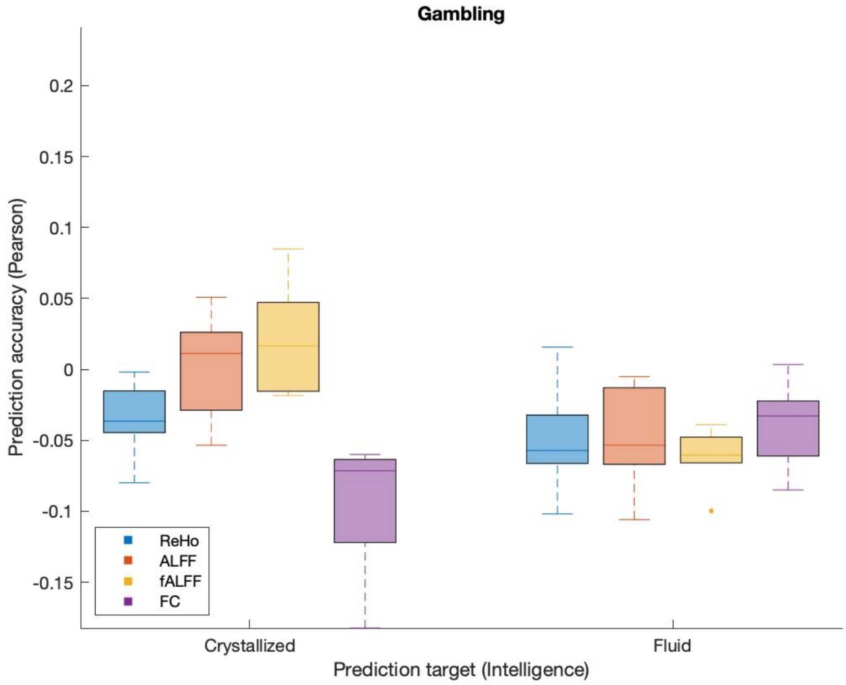

**Fig. S10:** Prediction of crystallized and fluid intelligence using the fingerprints of the Gambling fMRI task of the Human Connectome Project sample. Each boxplot shows the prediction performance across five cross-validation runs.

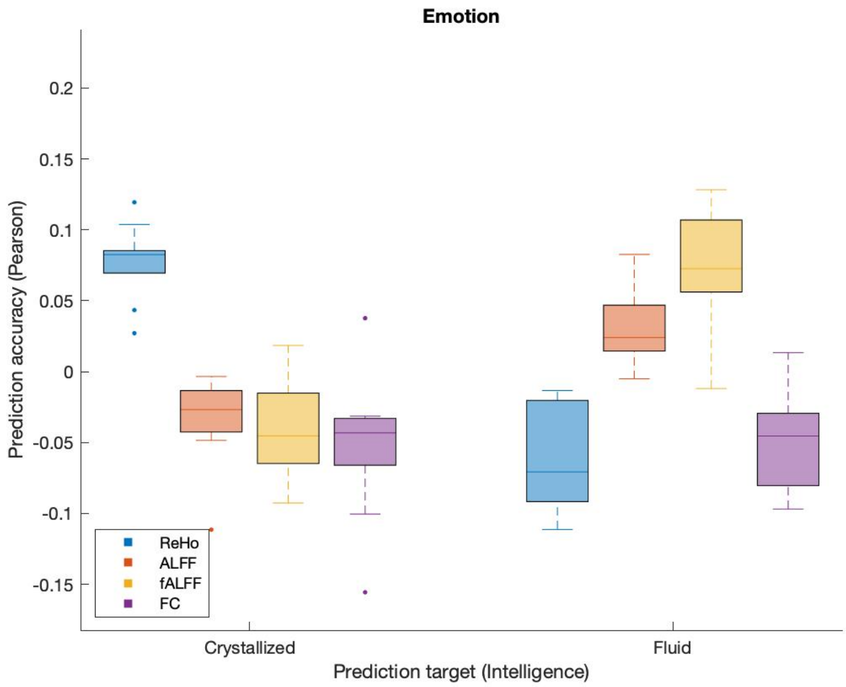


**Fig. S11:** Prediction of crystallized and fluid intelligence using the fingerprints of the Emotion fMRI task of the Human Connectome Project sample. Each boxplot shows the prediction performance across five cross-validation runs.

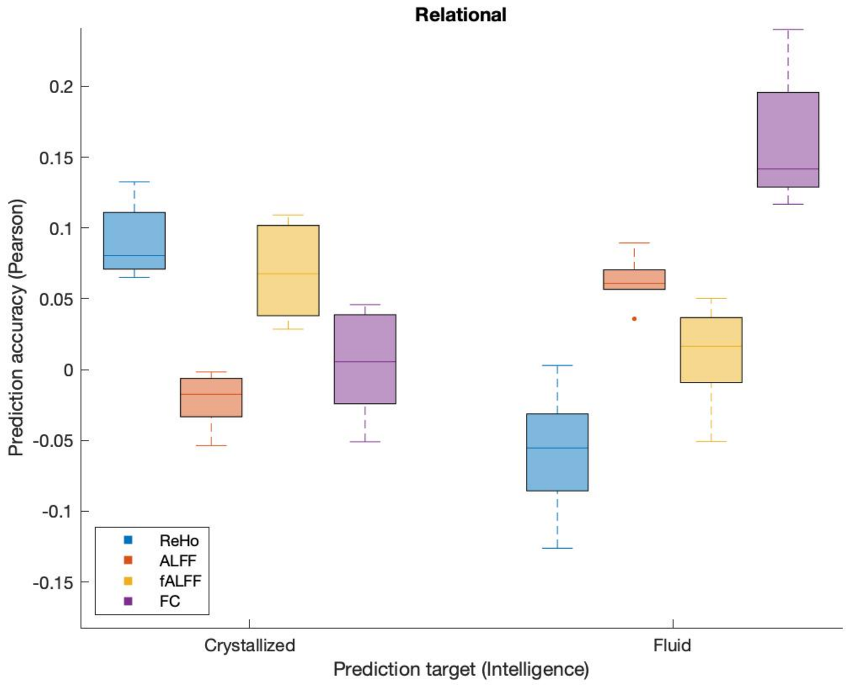

**Fig. S12:** Prediction of crystallized and fluid intelligence using the fingerprints of the Relational Processing fMRI task of the Human Connectome Project sample. Each boxplot shows the prediction performance across five cross-validation runs.

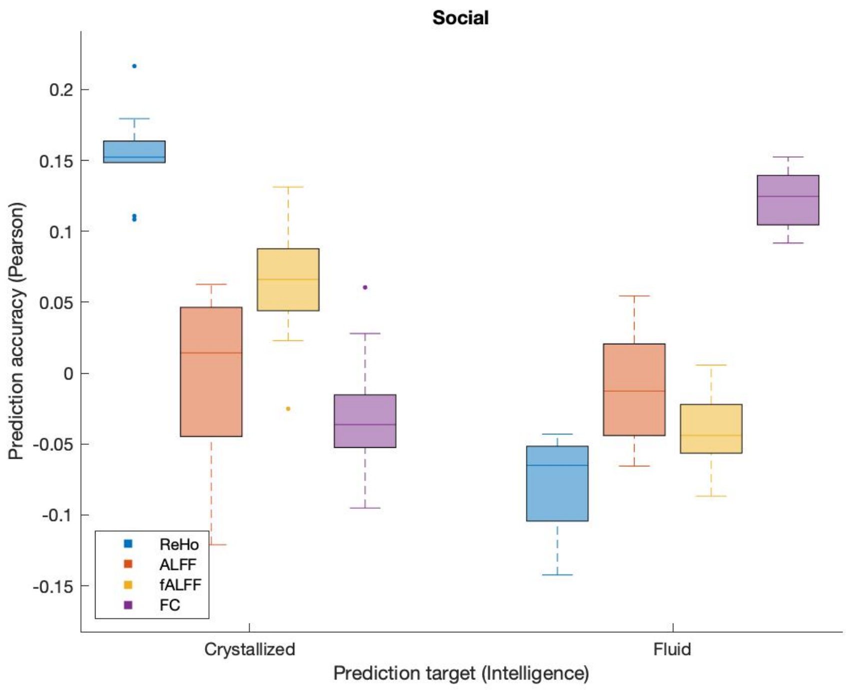


**Fig. S13:** Prediction of crystallized and fluid intelligence using the fingerprints of the Social Cognition fMRI task of the Human Connectome Project sample. Each boxplot shows the prediction performance across five cross-validation runs.

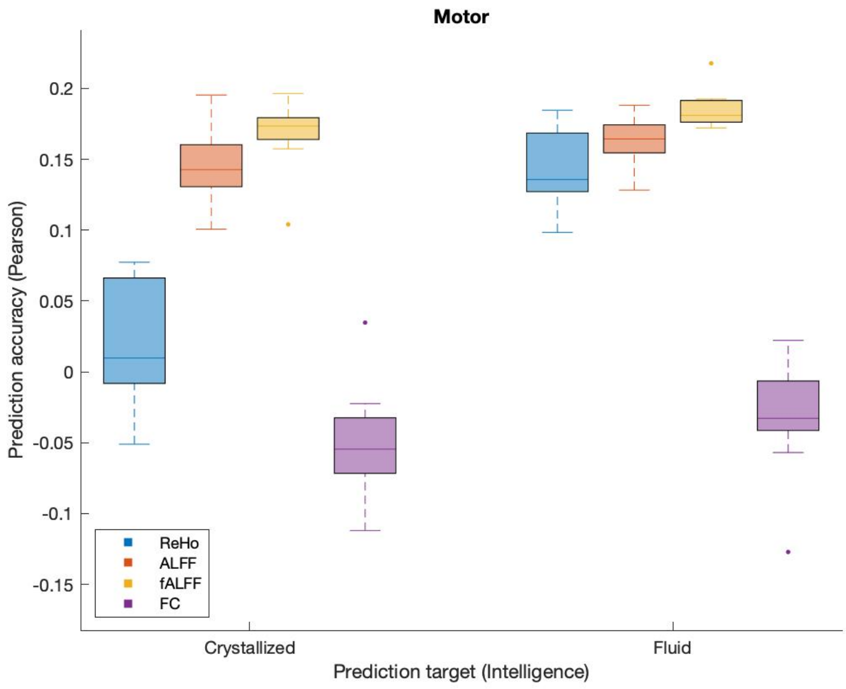


**Fig. S14:** Prediction of crystallized and fluid intelligence using the fingerprints of the Motor fMRI task of the Human Connectome Project sample. Each boxplot shows the prediction performance across five cross-validation runs.

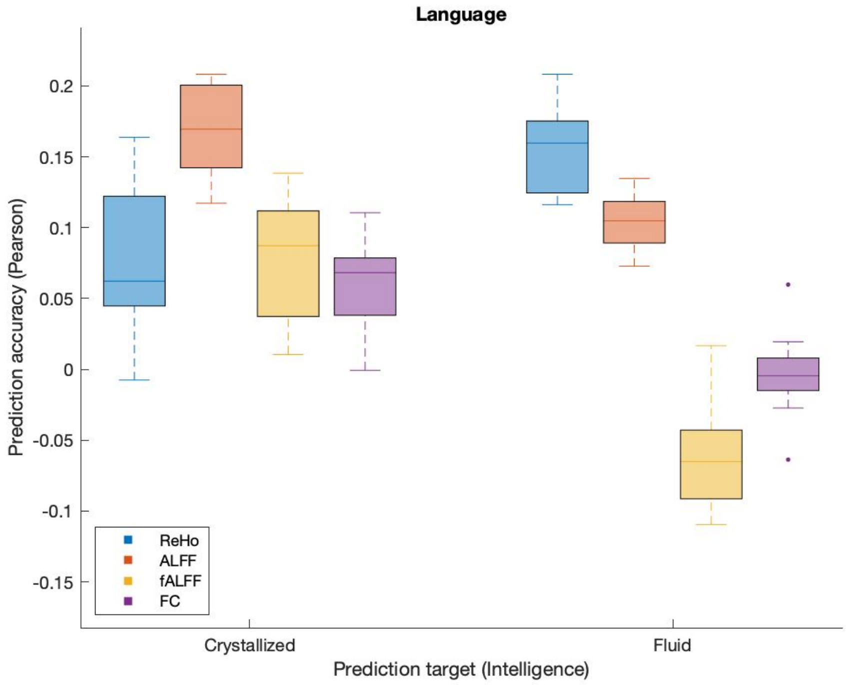


**Fig. S15:** Prediction of crystallized and fluid intelligence using the fingerprints of the Language fMRI task of the Human Connectome Project sample. Each boxplot shows the prediction performance across five cross-validation runs.

### Contribution of regions to individual uniqueness via *margin optimization*

The two optimization problems RS1-RS2 and RS2-Motor showed similar results with significantly increased margin (Fig. S7).

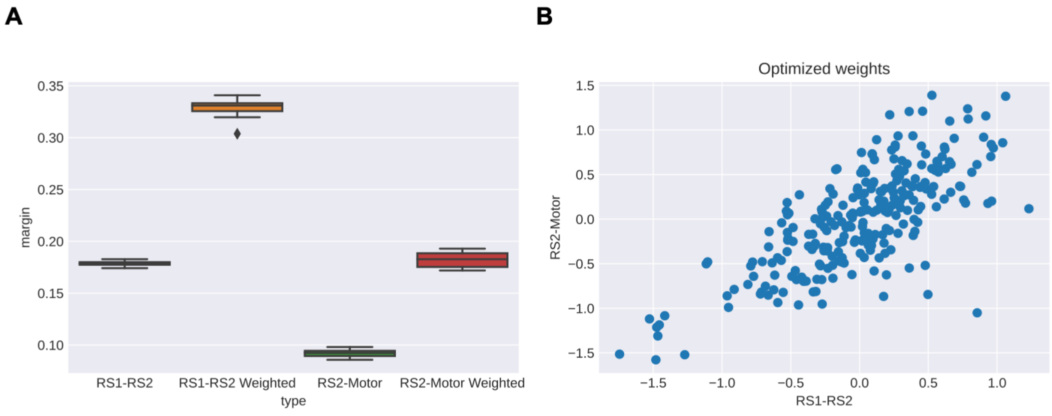


**Fig. S16**: **A)** improvement in the margin on the 50% out-of-sample subjects when using the weights derived on 50% of the data. The procedure was repeated 100 times with random sampling of train-test splits. The margin increased substantially. **B)** the averaged weights for the two optimization problems (RS1-RS2 and RS2-Motor) were highly correlated.

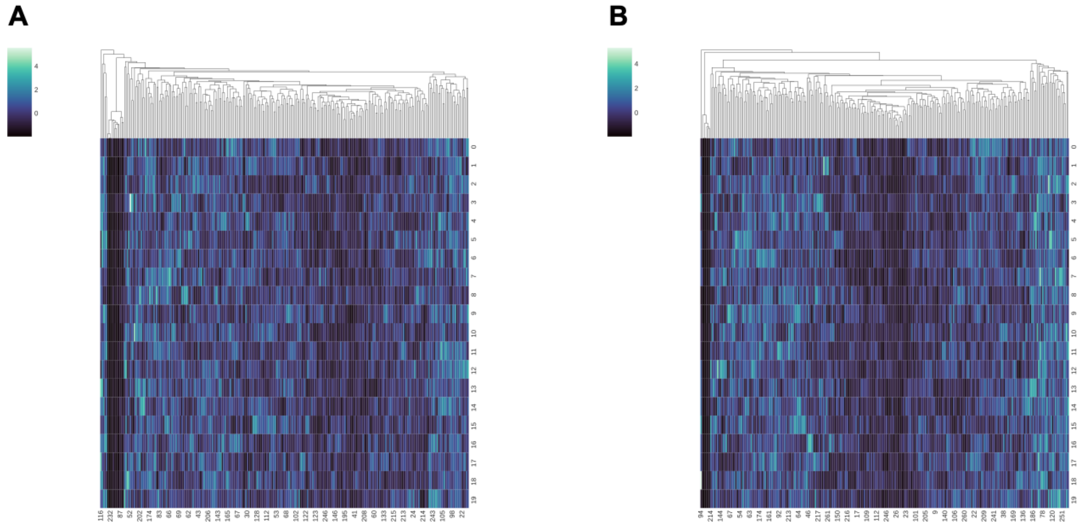

**Fig S17:** Weights across 100 runs of the optimization algorithm. The rows are the runs and the columns are the ROIs. **A)** RS1-RS2 runs. **B)** RS2-Motor runs. The weights were stable across the 100 optimization runs, therefore, for further analyses, we used the average of the weights across the 100 runs.

#### Assignment of ROIs to networks Yeo

**Table S1:** Proportion of the Power 3-mm ROI assignment to seven Yeo networks.

| None | Visual | Somato motor | Dorsal Attention | Ventral Attention | Limbic | Frontoparietal | Default Mode |
|---|---|---|---|---|---|---|---|
| 0.1 | 0.14 | 0.17 | 0.1 | 0.09 | 0.04 | 0.13 | 0.23 |

#### Correlations between intelligence and fingerprint stability in two clinical samples

**Table S2**: Correlations between subtests of fluid intelligence (Matrix Reasoning; 'WAIS_MATRICS_SCALE') or crystallized intelligence (Vocabulary; 'WAIS_VOCAB_SCALED') and fingerprint stability in the OpenNeuro.ds000115 sample.

| | Healthy controls | | | | Schizophrenia | | | |
|---|---|---|---|---|---|---|---|---|
| | ReHo | ALFF | fALFF | FC | ReHo | ALFF | fALFF | FC |
| Matrix Reasoning | r=0.10, p=0.68 | r=0.08, p=0.76 | r=0.30, p=0.22 | r=0.02, p=0.94 | **r=-0.50, p=0.03*** | **r=-0.65, p=0.002*** | r=-0.16, p=0.49 | **r=-0.60, p=0.005*** |
| Vocabulary | r=0.19, p=0.45 | **r=0.48, p=0.04*** | r=-0.02, p=0.94 | r=0.16, p=0.51 | r=0.06, p=0.80 | r=-0.22, p=0.35 | r=0.04, p=0.86 | r=-0.03, p=0.90 |

**Table S3**: Correlations between fluid cognition (composite score; 'nih_fluidcogcomp_unadjusted') or crystallized cognition (composite score; 'nih_crycogcomp_unadjusted'), and fingerprint stability in HCP Early Psychosis sample.

| | Healthy controls | | | | Non-affective psychosis | | | |
|---|---|---|---|---|---|---|---|---|
| | ReHo | ALFF | fALFF | FC | ReHo | ALFF | fALFF | FC |
| Fluid cognition | r=0.02, p=0.88 | r=-0.02, p=0.87 | r=0.01, p=0.93 | r=-0.01, p=0.95 | r=-0.03, p=0.79 | r=-0.04, p=0.77 | r=0.002, p=0.99 | r=0.09, p=0.46 |
| Crystallized cognition | r=-0.06, p=0.68 | r=-0.01, p=0.96 | r=0.18, p=0.22 | r=0.11, p=0.44 | r=-0.02, p=0.88 | r=0.01, p=0.91 | r=0.06, p=0.65 | r=0.16, p=0.18 |

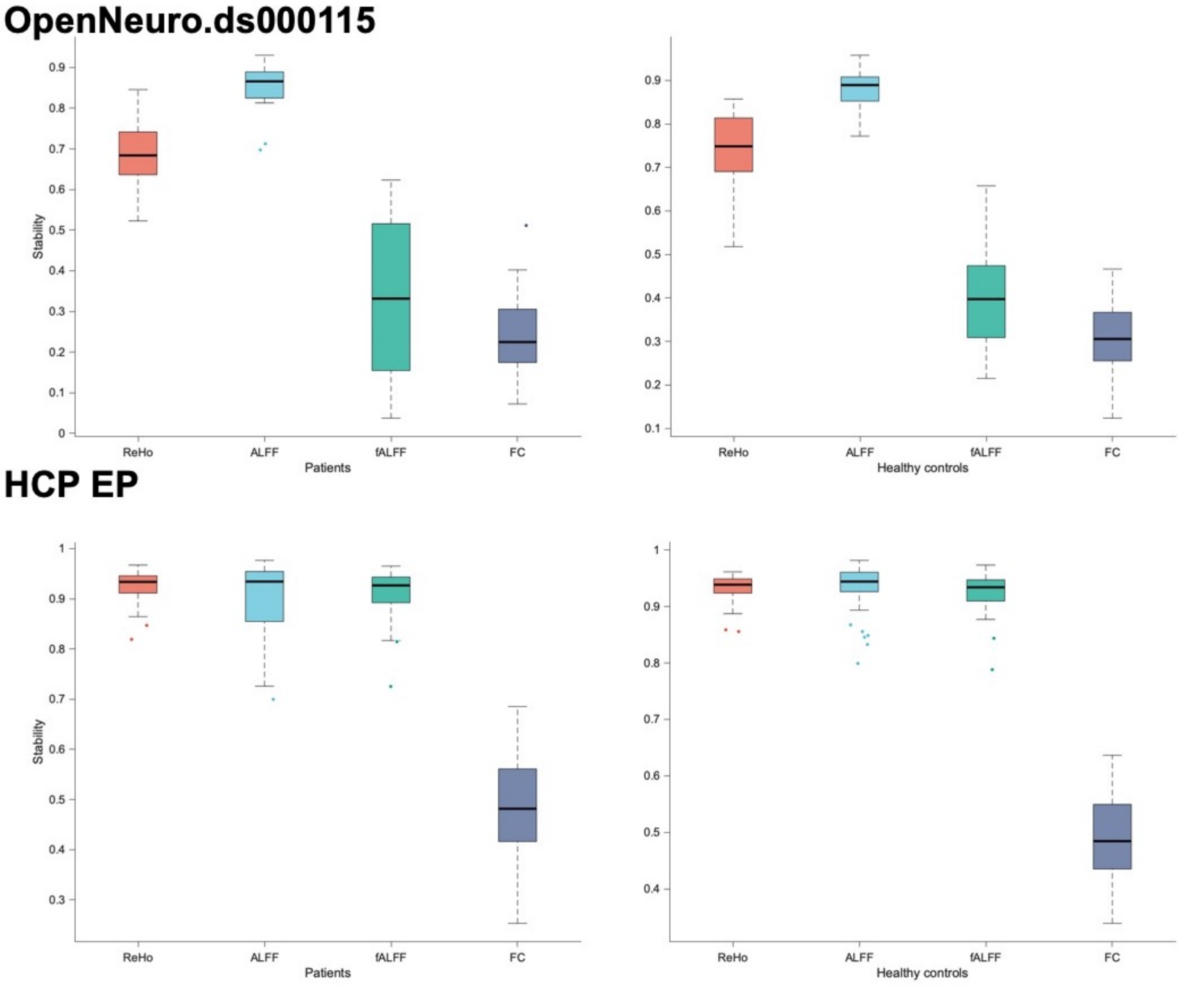


**Figure S18**: stability of fingerprints. Abbreviations: HCP EP = Human Connectome Project Early Psychosis sample.

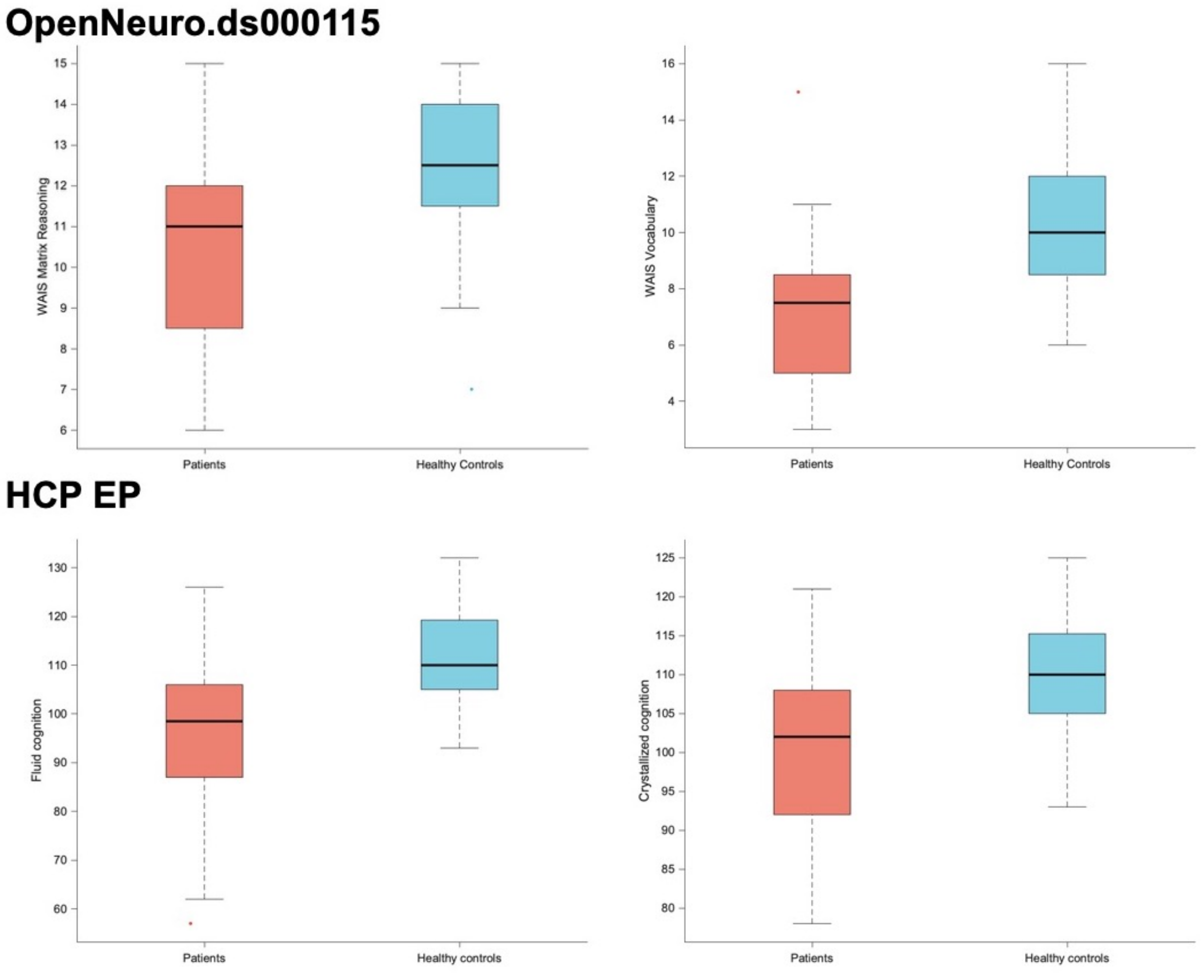


**Figure S19**: variation in intelligence scores.

### References


Chang, W., 2014. extrafont: Tools for using fonts.

Esteban, O., Markiewicz, C.J., Blair, R.W., Moodie, C.A., Isik, A.I., Erramuzpe, A., Kent, J.D., Goncalves, M., DuPre, E., Snyder, M., Oya, H., Ghosh, S.S., Wright, J., Durnez, J., Poldrack, R.A., Gorgolewski, K.J., 2019. fMRIPrep: a robust preprocessing pipeline for functional MRI. Nat. Methods 16, 111–116. doi:10.1038/s41592-018-0235-4

Finn, E.S., Shen, X., Scheinost, D., Rosenberg, M.D., Huang, J., Chun, M.M., Papademetris, X., Constable, R.T., 2015. Functional connectome fingerprinting: identifying individuals using patterns of brain connectivity. Nat. Neurosci. 18, 1664–1671. doi:10.1038/nn.4135

Glasser, M.F., Sotiropoulos, S.N., Wilson, J.A., Coalson, T.S., Fischl, B., Andersson, J.L., Xu, J., Jbabdi, S., Webster, M., Polimeni, J.R., Van Essen, D.C., Jenkinson, M., WU-Minn HCP Consortium, 2013. The minimal preprocessing pipelines for the Human Connectome Project. Neuroimage 80, 105–124. doi:10.1016/j.neuroimage.2013.04.127

Nooner, K.B., Colcombe, S.J., Tobe, R.H., Mennes, M., Benedict, M.M., Moreno, A.L., Panek, L.J., Brown, S., Zavitz, S.T., Li, Q., Sikka, S., Gutman, D., Bangaru, S., Schlachter, R.T., Kamiel, S.M., Anwar, A.R., Hinz, C.M., Kaplan, M.S., Rachlin, A.B., Adelsberg, S., Milham, M.P., 2012. The NKI-Rockland Sample: A Model for Accelerating the Pace of Discovery Science in Psychiatry. Front. Neurosci. 6, 152. doi:10.3389/fnins.2012.00152

Repovš, G., Barch, D.M., 2012. Working memory related brain network connectivity in individuals with schizophrenia and their siblings. Front. Hum. Neurosci. 6, 137. doi:10.3389/fnhum.2012.00137

Shen, X., Tokoglu, F., Papademetris, X., Constable, R.T., 2013. Groupwise whole-brain parcellation from resting-state fMRI data for network node identification. Neuroimage 82, 403–415. doi:10.1016/j.neuroimage.2013.05.081

Venkatesh, M., Jaja, J., Pessoa, L., 2020. Comparing functional connectivity matrices: A geometry-aware approach applied to participant identification. Neuroimage 207, 116398. doi:10.1016/j.neuroimage.2019.116398

Wickham, H., 2007. Reshaping Data with the reshape Package. J. Stat. Softw. 21. doi:10.18637/jss.v021.i12

Wickham, H., 2016. ggplot2: Elegant Graphics for Data Analysis. Springer-Verlag New York.

Zang, Y., Jiang, T., Lu, Y., He, Y., Tian, L., 2004. Regional homogeneity approach to fMRI data analysis. Neuroimage 22, 394–400. doi:10.1016/j.neuroimage.2003.12.030
